# Far-red light increases maize volatile emissions in response to volatile cues from neighboring plants

**DOI:** 10.1101/2022.09.12.507519

**Authors:** Rocío Escobar-Bravo, Bernardus C.J. Schimmel, Yaqin Zhang, Lei Wang, Christelle A.M. Robert, Gaétan Glauser, Carlos L. Ballaré, Matthias Erb

## Abstract

Plants perceive the presence and defense status of their neighbors through light and volatile cues, but how plants integrate both stimuli is poorly understood. We investigated if and how low Red to Far red light (R:FR) ratios, indicative of shading or canopy closure, affect maize (*Zea mays*) responses to herbivore-induced plant volatiles (HIPVs), including the green leaf volatile (*Z*)-3-hexenyl acetate. We modulated light signaling and perception by using FR supplementation and a *phyB1phyB2* mutant, and we determined volatile release as a response readout. To gain mechanistic insights, we examined expression of volatile biosynthesis genes, hormone accumulation, and photosynthesis. Exposure to a full blend of HIPVs or (*Z*)-3-hexenyl acetate induced maize volatile release. Short-term FR supplementation increased this response. In contrast, prolonged FR supplementation or constitutive phytochrome B inactivation in *phyB1phyB2* plants showed the opposite response. Short-term FR supplementation enhanced photosynthesis and stomatal conductance and (*Z*)-3-hexenyl acetate-induced JA-Ile levels. We conclude that a FR-enriched light environment can prompt maize plants to respond more strongly to HIPVs emitted by neighbors, which might be explained by changes in photosynthetic processes and phytochrome B signaling. Our findings reveal interactive responses to light and volatile cues with potentially important consequences for plant-plant and plant-herbivore interactions.

## Introduction

The capacity to perceive and respond to fluctuating environments is essential to life on earth. As primary producers in terrestrial ecosystems, plants are constantly dealing with competitors, pests, and pathogens. Consequently, they have evolved systems to detect these stressors and respond to them appropriately. While recent work has advanced our knowledge of how plants perceive individual environmental cues to detect biotic stressors, much less is known about their capacity to simultaneouly integrate multiple ones (Nguyen *et al*., 2016; Escobar-Bravo *et al*., 2017).

Plants can perceive qualitative and quantitative changes in light. When plants grow in dense canopies there is a strong decline in photosynthetically active radiation (PAR) and a change in light quality due to the light absorption and reflection by leaves (Ballaré & Pierik, 2017). Plants typically absorb red (R) and blue (B) light whereas the far red (FR) component of the sunlight is mostly reflected and transmitted. Consequently, in closing or dense canopies, there is a decrease in the R to FR ratio (R:FR) resulting from the reduction in R light, which is strongly absorbed by chlorophylls, and the increase in FR radiation. A reduction in R:FR ratios is a signal of foliar shading and proximity of other plants (Ballaré *et al*., 1990). Plants can respond to this FR-enriched light environment by upward bending of the leaves (hyponasty) and stem elongation, which allows them to reach more illuminated areas and outcompete their neighbors (Franklin, 2008; Ballaré & Pierik, 2017). Variations in R:FR ratios are perceived by the photoreceptor phytochrome B (phyB) (Fankhauser, 2001). PhyB is inactivated under low R:FR, which triggers the expression of growth-related pathways controlled by hormones such as auxins and gibberellins (Fiorucci & Fankhauser, 2017).

Besides light cues, plants can perceive and respond to volatile organic compounds emitted by herbivores (Helms *et al*., 2017) and neighboring plants (Hu, 2022). When a plant is attacked, it releases distinct blends of herbivore induced plant volatiles (HIPVs). HIPVs can be perceived by non-attacked plants and induce or prime their defenses (Engelberth *et al*., 2004). Plant responses to HIPVs includes the rapid release of volatiles (Engelberth *et al*., 2004) and HIPV-induced priming, which allows plants to respond more rapidly and strongly once they are under attack (Ton *et al*., 2007). In maize, HIPVs such as indole and (*Z*)-3-hexenyl acetate (HAC) prime the jasmonic acid (JA) pathway (Erb *et al*., 2015; Ye *et al*., 2019), triggering the deployment of an array of defenses that enhance herbivore resistance and the attraction of parasitoids (Ton *et al*., 2007). In addition, HAC can directly induce defenses, and volatile biosynthesis and release (Engelberth *et al*., 2004; Hu *et al*., 2019).

Recent work has demonstrated that R:FR ratios influence defense deployment in plants, including volatiles. Reduced R:FR via FR supplementation to ambient light suppresses direct defenses against herbivores and pathogens (Moreno *et al*., 2009; Cargnel *et al*., 2014; Courbier *et al*., 2021). This is predominantly mediated by crosstalk between phyB and the JA pathway (Moreno *et al*., 2009; Campos *et al*., 2016; Fernández-Milmanda *et al*., 2020). When phyB is inactivated, JA signaling is attenuated as a consequence of reduced levels of bioactive JA forms (Fernández-Milmanda *et al*., 2020), decreased stability of MYC transcription factors (Chico *et al*., 2014), and enhanced stability of JAZ transcriptional suppressors (Chico *et al*., 2014; Leone *et al*., 2014). As volatile biosynthesis can be regulated by JA, this might explain why phyB inactivation by low R:FR ratios modulates the emission of some volatile compounds. Upon FR supplementation, for instance, barley (*Hordeum vulgare*) plants show reduced constitutive emission of several terpenes but an increased emission of the monoterpene linalool oxide (Kegge *et al*., 2015). Similarly, reduced R:FR via FR supplementation in combination with shading (i.e., low PAR) reduces constitutive and methyl jasmonate-induced terpenes and green leaf volatiles emissions in *Arabidopsis thaliana*, but induces the emission of ethylene, a volatile phytohormone commonly deployed upon low R:FR (Kegge *et al*., 2013). The fact that emission of specific volatiles is enhanced under low R:FR (Kegge *et al*., 2015; Cortés *et al*., 2016) suggests that the response to competition signals may be highly specific.

Unlike the well documented effects of low R:FR ratios on plant defenses and volatile emissions, it is unknown if FR can affect plant’s perception and responses to volatile cues emitted by neighbors. That plants are capable of sensing and integrating different types of information is illustrated, for instance, by seed germination initiation, which requires a combination of environmental cues such as temperature, moisture, and light (Finch-Savage & Leubner-Metzger, 2006). Furthermore, plants that are under attack by herbivores of different feeding guilds often show unique response patterns due to the hormonal integration of different signals (Soler *et al*., 2013; Zhang *et al*., 2013; Kroes *et al*., 2015; Chrétien *et al*., 2018). Thus, plants might also integrate light and volatiles cues into different responses. For instance, they might reduce their response to HIPVs when they perceive light cues that indicate the close proximity of neighboring plants as it may reduce the risk of an individual plant being attacked (Barbosa *et al*., 2009).

In this study, we have investigated the following research questions: (1) Does FR enriched light environment modulate maize volatile emissions in response to HIPVs or the green leaf volatile HAC? If so, (2) How does FR enriched light environment modulate maize volatile emissions in response to the volatile cue HAC? and (3) Is FR-mediated changes in volatile emissions mediated by the inactivation of the PhyB photoreceptor in maize?

To investigate if and how maize (*Zea mays*) plants integrate light and volatile cues from neighbors into distinct volatile emissions, we employed a newly developed automated high-throughput volatile screening platform with a programmable multi-channel LED lighting and a Vocus PTR-TOF-MS system. We exposed maize plants to short-term FR supplementation as a light cue for vegetation shade and to HIPVs or HAC alone as volatile cues for the presence of herbivore-infested neighbors, and we determined direct induction and priming of volatile emissions in time-series analyses. To gain mechanistic insights, we determined gene expression, phytohormone levels, and photosynthetic responses in response to short-term FR treatments and HAC. Finally, we used a maize *phyB_1_phyB*_2_ double mutant and short-or long-term FR supplementation experiments to determine the role of long-term inactivation of PhyB in FR-induced volatile emissions in response to HAC. Our experiments demonstrate that short-term exposure to a FR-enriched light environment enhances maize volatile emissions in response to HIPVs or HAC alone, illustrating how plants use different types of signals from neighbors to reprogram their own phenotypes.

## Materials and Methods

### Plant material

Experiments were conducted with the maize (*Zea mays*) inbred line B73, the *phyB_1_phyB*_2_ maize double mutant, and its wild-type W22. The *phyB_1_phyB*_2_ double mutant was provided by Prof. Michael Scanlon (School of Integrative Plant Science, Cornell University). This mutant is homozygous for the Mutator (Mu) transposon insertion alleles *phyb1-563* and *phyb2-12058* resulting in no detectable PHYB protein (Sheehan *et al*., 2007; Dubois *et al*., 2010). B73 responds to HAC (Hu *et al*., 2018) and FR (Dubois *et al*., 2010) and releases moderate volatile emissions upon herbivory (Block *et al*., 2018). For experiments with 11-day-old plants, seeds were sown in transparent cylindric plastic pots (11×4 cm) filled with commercial soil (Selmaterra, BiglerSamen, Switzerland) and wrapped with aluminum foil to prevent root exposure to light. For experiments with 30-day-old plants, seeds were sown in black plastic pots (9×9×10 cm) filled with the same commercial soil. Plants were grown in a ventilated greenhouse supplemented with artificial lighting under 50-70% relative humidity, 14/10 h light/dark photoperiod, and 14-22/10-14°C day/night temperatures.

### Insects

Eggs of *Spodoptera littoralis* were provided by Oliver Kindler (Syngenta, Stein, CHE) and the larvae were reared on artificial diet as described in Maag *et al*., 2014. *S. littoralis* oral secretions were collected from third-to fourth-instar larvae as described in Hu *et al*., 2019 and stored at − 80°C until use.

### Light treatments

Light treatments were conducted in a custom-made high-throughput phenotyping platform designed for volatile analyses. The platform consisted of 96 transparent cylindrical glass chambers (120 mm Ø, 450 mm height; abon life sciences, Switzerland) closed at the top with transparent glass lids and distributed over two tables. Each table was supplied with four light emitting diodes (LEDs) lamps placed above the glass chambers. Each lamp contained 120 LEDs with nine wavebands consisting of 380, 400, 420, 450, 520, 630, 660, 735 nm and 5700K white (RX30, Heliospectra). Individual glass chambers were supplied with an airflow of 0.8 L/min and an air outlet. Plants were placed inside these glass chambers and exposed to low (∼0.5) or high (∼2.65) R:FR ratios (R, λ = 600-700 nm; FR, λ = 700-800 nm) by providing supplemental FR to the same white light background (120 ± 15 μmol m^-2^ s^-1^) (Supplemental Fig. S1) with a photoperiod of 16/8 h light/dark. This low R:FR corresponds to the ratios observed in high density maize canopies (Maddonni *et al*., 2002), where low PAR levels as the ones used in our experimental set-up are commonly observed (Xue *et al*., 2016). Light treatments were separated by white opaque curtains and started between 9:00 and 10:00 AM two days before the volatile exposure treatments.

#### (Z)-3-hexenyl acetate treatment

Individual plants were exposed to one HAC dispenser for 6 h within the transparent cylindrical glass chambers of the custom-made high-throughput phenotyping platform described above. HAC dispensers were prepared as described in Hu *et al*., 2019, each releasing approximately 70 ng h^−1^, which has been previously described in herbivore-attacked maize plants (Mérey *et al*., 2011). HAC treatments started between 9:00-10:00 AM.

#### Simulated herbivory

Two leaves (leaf 2 and 3 from the bottom) per plant were wounded over an area of ca. 0.5 cm^2^ parallel to the central vein using a haemostat followed by the application of 10 μL of *S. littoralis* larval oral secretions (diluted 1:1 in autoclaved distilled water) in each wounding site. This treatment induces plant defense responses comparable to real herbivory (Erb *et al*., 2009).

#### Linalool dispensers

Linalool dispensers were prepared as described in Hu *et al*., 2019 for HAC but with some modifications. They contained 100 µL of pure linalool (SIGMA) and were pierced through the rubber septum with a 1 µL micropipette (Drummond, Millan SA, Switzerland).

#### PTR-TOF-MS volatile analysis

Plant volatile emission was determined using the automated high-throughput phenotyping platform described above for light treatments. Volatiles emitted by a plant inside each transparent glass chamber were purged by a clean air flow (0.8 L/min) that carried the emitted volatiles directly to a Vocus proton transfer reaction time-of-flight mass spectrometer (PTR-TOF-MS) system (Tofwerk, Switzerland) for analysis. The Vocus PTR-TOF-MS was operated using H_3_O^+^ as the reagent ion and was integrated into an autosampler (Gonin *et al*., 2022) that cycles between individual chambers/plants. Each plant was measured for 50 or 60 seconds. Between individual plants, a fast zero air measurement was performed for 5 seconds. Complete mass spectra (0-500 Th) were saved and plotted at 5 Hz (5 unique mass spectra per second). PTR-TOF-MS data processing consisted of mass calibration and calculation of peak intensities using the Tofware software package v3.2.0, followed by normalization of the signals to aboveground (stem and leaves) plant fresh biomass.

#### GC-MS volatile emission analysis

Dynamic volatile collection was conducted using adsorbent Super-Q traps followed by gas chromatography-mass spectrometry (GC-MS) analysis of volatile extracts. For this, air was pulled out from the PTR-TOF-MS transparent glass chambers containing the test plants through the air outlet ports (0.5 L/min) and across the Super-Q traps (Porapak-Q^TM^ 50 mg, Volatile collection Trap, USA). Volatiles were collected for periods of 2 h for 12 h. After each collection period, Super-Q traps were exchanged for new ones and the volatiles collected on the trapping filters were extracted with 200 μl of Dichloromethane (Roth) containing 200 ng of both N-octane and nonyl-acetate as internal standards. Extracts were analyzed by GC-MS (Agilent 7820A GC interfaced with and Agilent 5977E MSD, Palo Alto, CA, USA). Aliquots of 1 μL were injected into the column (HP5-MS, 30 m, 250 μm ID, 2.5-μm film, Agilent Technologies, Palo Alto, CA, USA) using a pulsed split-less mode and He as carrier gas at a flow rate of 1 mL min^-1^. A temperature gradient of 5°C min ^-1^ from 40°C to 200°C was used. Compound identification was based on similarity to library matches (NIST search 2.2 Mass Spectral Library. Gaithersburg, MD, USA), retention times, and spectral comparison with pure compounds. Relative abundance was calculated by dividing the normalized peak area (peak area analyte/peak area of the internal standard N-octane) of individual compounds by aboveground plant fresh biomass.

#### Phytohormone analysis

The concentrations of 12-oxo-phytodienoic acid (OPDA), JA, jasmonic acid-isoleucine (JA-Ile), salicylic acid (SA), abscisic acid (ABA) and auxin (indole acetic acid, IAA) were determined in aboveground tissues (stem and leaves) following the procedure described in Glauser *et al*., 2014 with minor modifications (Supplemental Methods S1).

#### Gene expression analysis

Total RNA was isolated using TRI Reagent (Sigma-Aldrich) following the manufacturer’s instructions. DNase treatment, reverse transcription, and first strand cDNA synthesis were performed using gDNA Eraser (Perfect Real Time) and PrimeScript Reverse Transcriptase following manufacturer’s instructions (Takara Bio Inc. Kusatsu, Japan). For gene expression analysis, 2 μL of cDNA solution (equivalent to 10 ng of total RNA) served as template in a 10 μL qRT-PCR reaction using ORA™ SEE qPCR Mix (Axonlab) on an Applied Biosystems® QuantStudio® 5 Real-Time PCR System. Normalized expression (NE) values were calculated as in Alba *et al*., (2015). Gene identifiers, primer sequences, and references are listed in Table S1.

#### Stomatal conductance, density, and net photosynthesis

Stomatal conductance was measured using an AP4 Porometer (Delta-T Devices Ltd) with cup aperture dimension of 2.5 x 17.5 mm. Stomatal density was determined in epidermal imprints of the abaxial surface using clear nail varnish. Epidermal imprints were imaged using a Leica DM2500 optical microscope coupled to a Leika HD Digital microscope camera (Leica MC170 HD) (Leica Microsystems, Wetzlar, Germany). The number of stomata was determined in 10 pictures per plant and leaf using ImageJ software. Net photosynthesis was determined using a LI-6400 Portable Photosynthesis system (LI-COR, Biosciences).

#### Statistical analysis

Data analysis was conducted in R version 4.0.4 (R core Team, 2016). Effects of (1) sampling time, (2) light, (3) genotype, (4) volatile exposure, and (5) their interactions on volatile emissions in the different experiments were tested in linear mixed-effects models (LME) using nlme library. We included individual plants as a random factor and a correlation structure when autocorrelation among residuals was found to be significant (*p* < 0.05) in the models. Fitted models were subjected to type III analyses of variance (ANOVAs) to produce a summary of the F-and p statistics (car package Fox & Weisberg, 2018). When needed, data from volatiles signatures were log10 or squared root transformed prior analysis to correct for heteroscedasticity. In Fig. 4 differences among light treatments were tested for all volatiles at each sampling time by estimated marginal means. Hormone concentrations and gene expression were analyzed separately for each time point by two-way ANOVAs followed by Tukey HSD post hoc test when light, HAC, or their interaction were significant. Effects of light, HAC, herbivory, and their interactions on stomatal conductance, density, and net photosynthesis were tested by two-way ANOVAs followed by Tukey HSD post hoc test.

**Figure 1.**
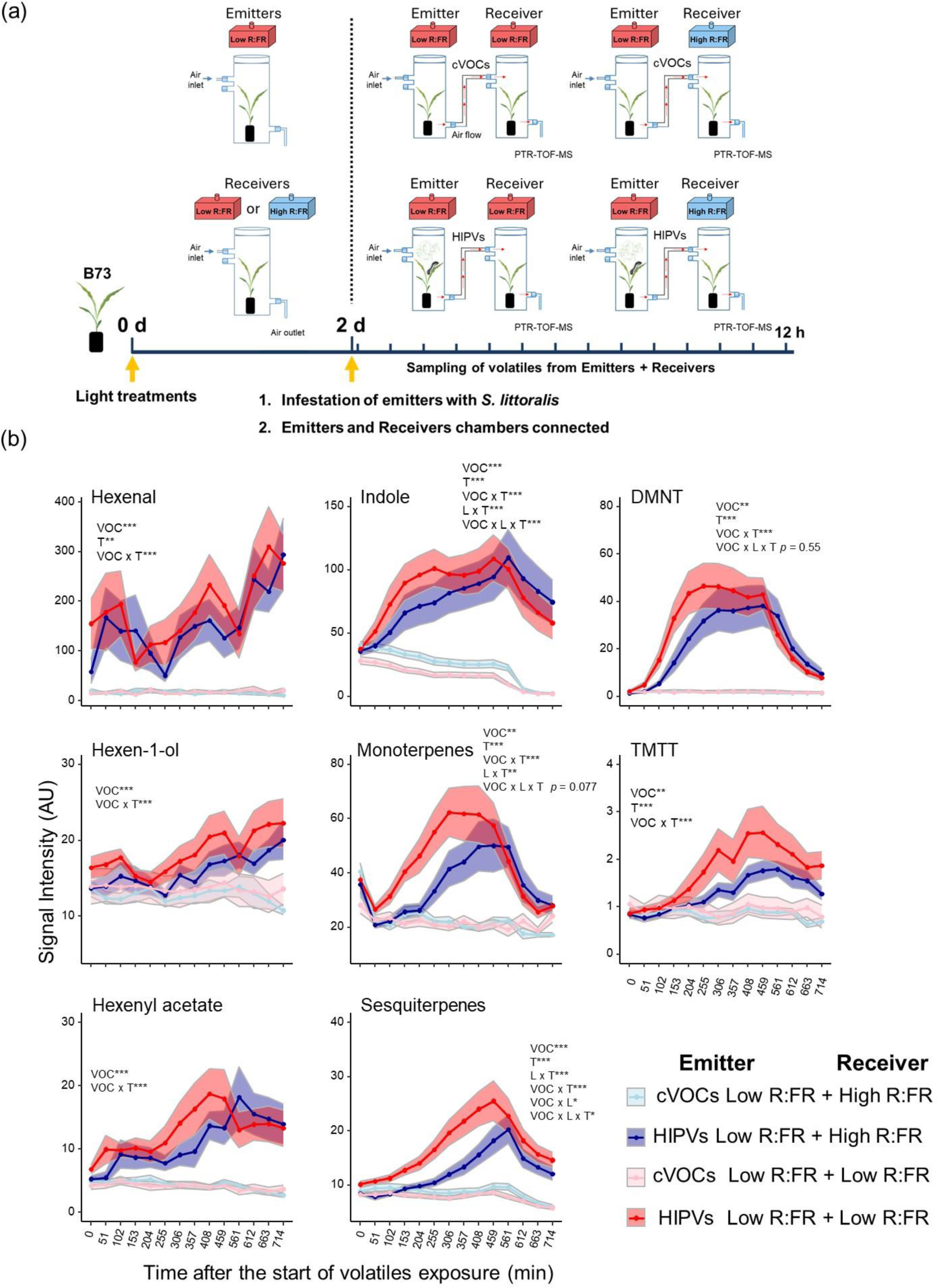
Far-red enrichment boosts maize responses to herbivore-induced plant volatiles. (a) Schematic overview of the experimental set-up. Eleven-day old ‘B73’ maize (*Zea mays*) plants were exposed to low or high R:FR light conditions by modulating FR levels. After two days, receiver plants growing under low or high R:FR were exposed to constitutive volatiles (cVOCs) or *S. littoralis* herbivory-induced plant volatiles (HIPVs) from emitter plants exposed to low R:FR. Emitter and Receiver plants were connected via Teflon tubing and their combined volatile emissions were measured by PTR-TOF-MS for 12 h. Volatile measurements started at 12:00 a.m. **(b)** Time-series analysis of volatile emissions (mean ± SEM, *n* = 12) collected from paired (Emitter + Receiver) plants. The effects of sampling time (T), light treatments of receivers (L), volatiles from emitters (VOC), and their interactions on volatile emissions were tested using linear mixed-effects models. Statistically significant effects are shown in each graph (\**p* < 0.05, ** *p* < 0.001, *** *p* < 0.001). DMNT and TMTT stand for the homoterpenes (*E*)-4,8-dimethyl-1,3,7-nonatriene and (*E*, *E*)-4,8,12-trimethyltrideca-1,3,7,11-tetraene, respectively. AU refers to arbitrary units.

**Figure 2.**
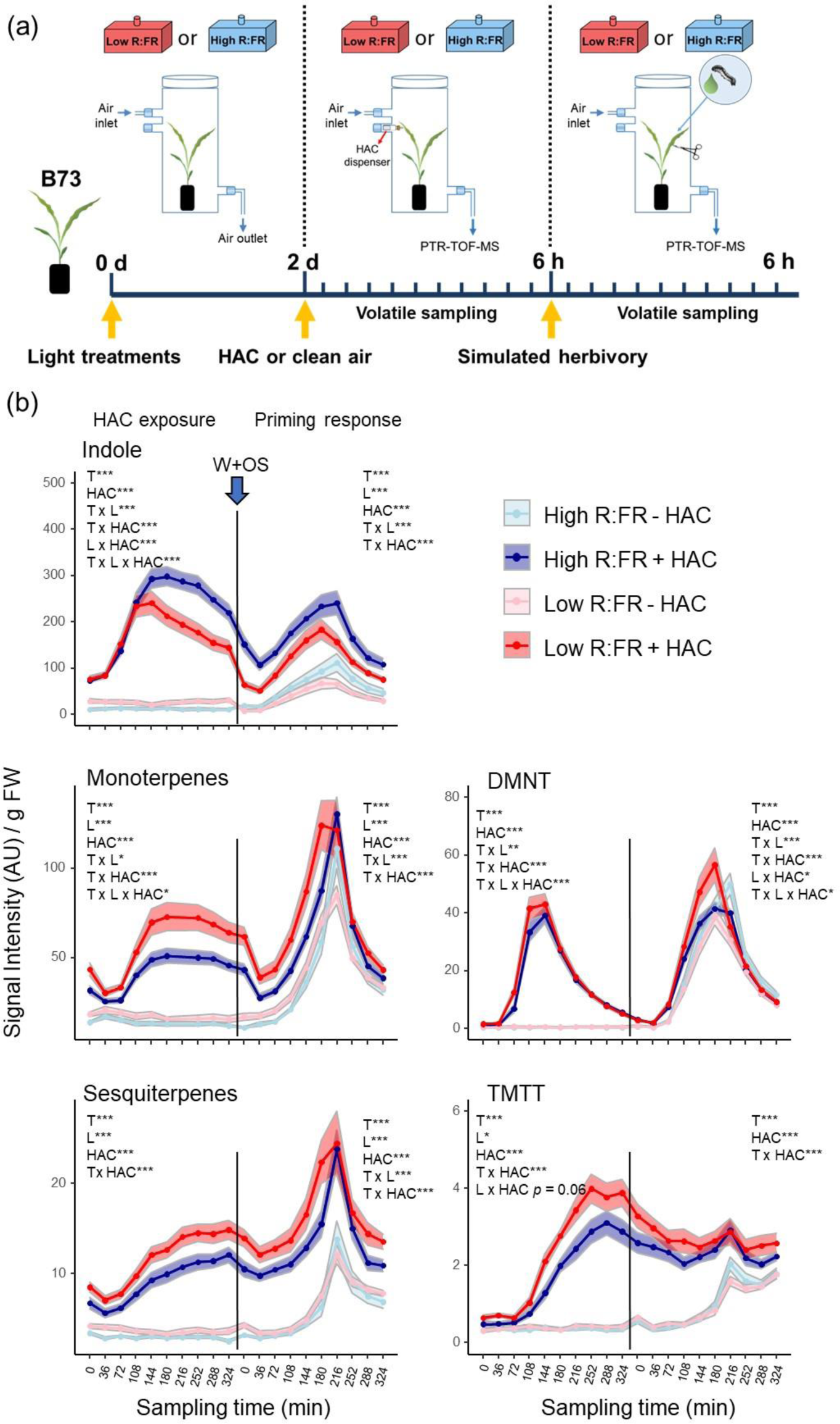
Far-red enrichment boosts maize responses to (*Z*)-3-hexenyl acetate. **(a)** Schematic overview of the experimental set-up. Eleven-day old ‘B73’ maize (*Zea mays*) plants were exposed to low or high R:FR conditions for 2 d by modulating FR levels, and subsequently exposed to (*Z*)-3-hexenyl acetate (HAC) or clean air for 6 h. After HAC or clean air treatment, all plants were induced with simulated herbivory (wounding and application of *Spodoptera littoralis* oral secretions; W+OS). Volatile emissions were determined during HAC exposure and after simulated herbivory by PTR-TOF-MS. **(b)** Time-series analysis of volatile emissions (mean ± SEM, *n* = 9) determined in low and high R:FR-treated plants during exposure to HAC (+ HAC) or clean air (-HAC) and after induction with simulated herbivory (priming phase). HAC exposure and volatile measurements started between 9:00 and 10:00 am. The effects of the sampling time (T), HAC, light treatment (L) and their interactions on volatile emissions were tested separately for the HAC exposure and priming phase using linear mixed-effects models. Statistically significant effects are shown in each graph (\**p* < 0.05, ** *p* < 0.001, *** *p* < 0.001). DMNT and TMTT stand for the homoterpenes (*E*)-4,8-dimethyl-1,3,7-nonatriene and (*E*,*E*)-4,8,12-trimethyltrideca-1,3,7,11-tetraene, respectively. AU refers to arbitrary units.

**Figure 3.**
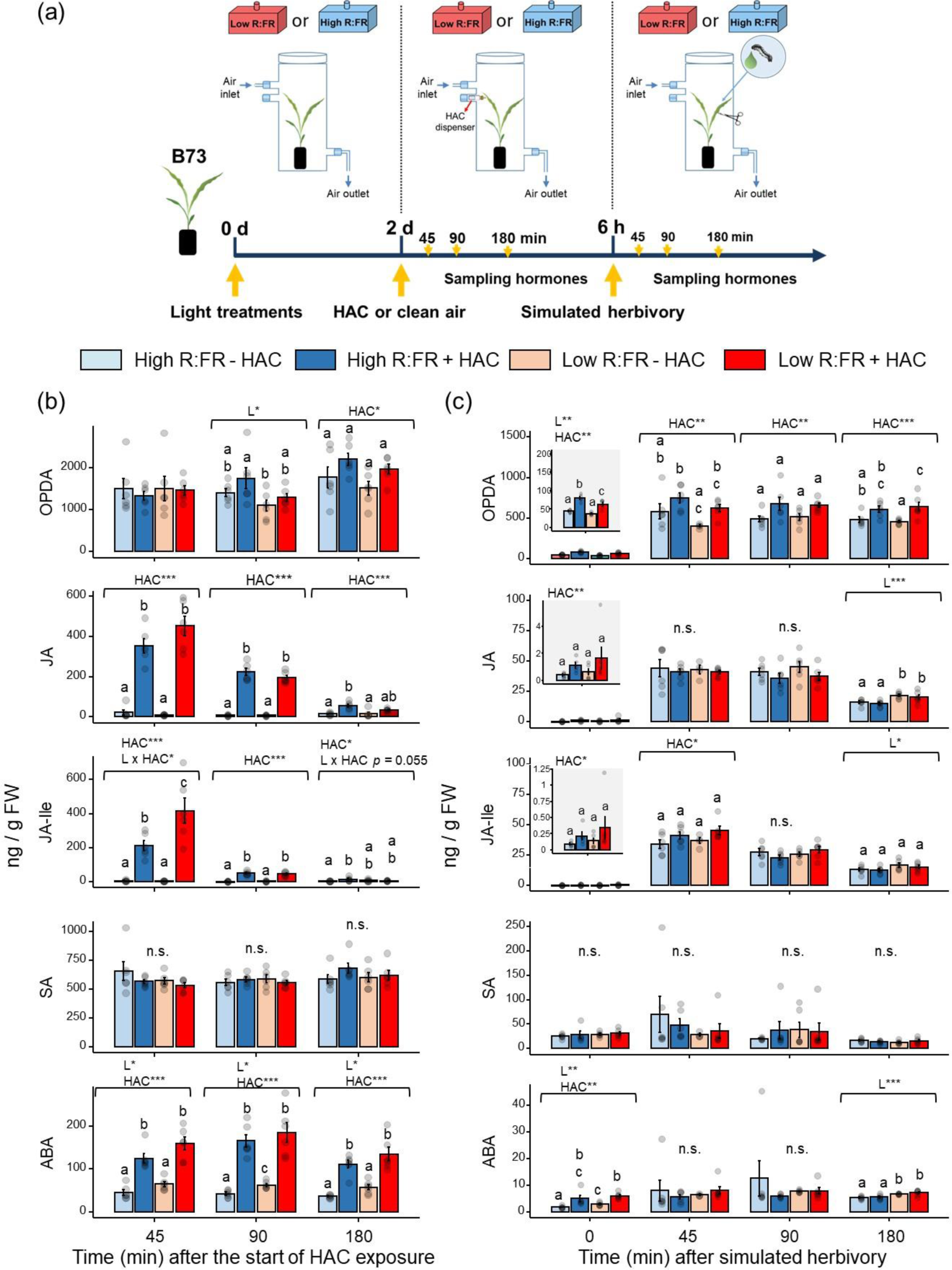
Far-red enrichment enhances JA-Ile accumulation in response to (*Z*)-3-hexenyl acetate. **(a)** Schematic overview of the experimental set up and sampling times. Concentrations (mean ± SEM, *n* = 4-6) of 12-oxo-phytodienoic acid (OPDA), jasmonic acid (JA), jasmonic acid-isoleucine (JA-Ile), salicylic acid (SA), and abscisic acid (ABA) in high and low R:FR treated B73 maize (*Zea mays*) plants that were in **(b)** exposed to (*Z*)-3-hexenyl acetate (+HAC) or clean air (-HAC) and in **(c)** all induced with simulated herbivory (wounding and application of *Spodoptera littoralis* oral secretions) after exposure to HAC or clean air for 6 h. Effects of light treatment (L), HAC exposure and their interaction were tested by two-way ANOVA at each time point. Different letters denote significant differences among groups tested by Tukey-HSD post hoc test. Statistically significant effects are shown in each graph (\**p* < 0.05, ** *p* < 0.001, *** *p* < 0.001).

**Figure 4.**
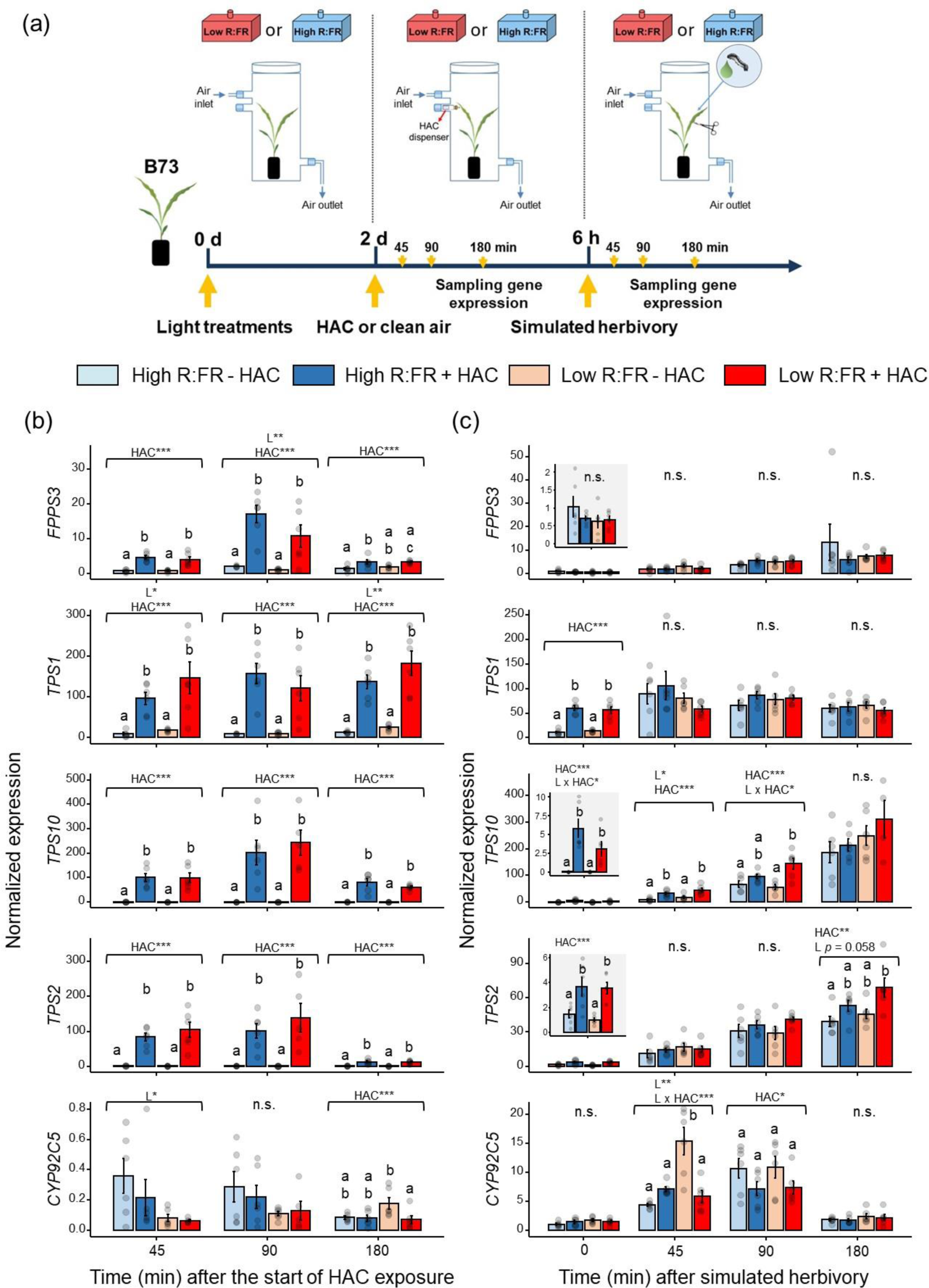
Far-red enrichment modulates the expression of volatile biosynthetic genes. **(a)** Schematic overview of the experimental set up and sampling times. Normalized expression (mean ± SEM, *n* = 6) of terpene metabolism-related genes determined in aboveground (leaves and stem) tissues of low and high R: FR-irradiated B73 maize (*Zea mays*) plants that were in **(b)** exposed to (*Z*)-3-hexenyl acetate (HAC) or clean air and in **(c)** all induced with simulated herbivory (wounding and application of *Spodoptera littoralis* oral secretions) after exposure to HAC for 6 h. Effects of light treatment (L), HAC and their interaction were tested by two-way ANOVA at each time point. Different letters denote significant differences among groups tested by Tukey-HSD post hoc test. Statistically significant effects are shown in each graph (\**p* < 0.05, ** *p* < 0.001, *** *p* < 0.001) Abbreviations for precursors and genes: *FPPS3*, farnesyl pyrophosphate synthase 3; *TPS1, Terpene synthase 1; TPS10*, *Terpene synthase 10*; *TPS2*, *terpene synthase 2*; *CYP92C5*, *P450 Monooxygenase CYP92C5*.

## Results

### Far-red enrichment boosts maize responses to herbivore-induced plant volatiles

Herbivore-induced plant volatiles can induce the emission of volatiles in non-attacked plants and enhance their defense responses when actual herbivory occurs (Farag & Paré, 2002; Engelberth *et al*., 2004). To determine whether different R:FR ratios alter responses to HIPVs in maize, we first exposed 11-day old B73 seedlings to high or low R:FR for two days by modulating FR levels and measured growth-related parameters (Supplemental Fig. S2). Low R:FR increased the elongation of stem internodes and auxin levels (Supplemental Fig. S2bc), confirming the effectiveness of our FR treatment in triggering shade avoidance (Dubois & Brutnell, 2009).

Next, we compared volatile emissions in high and low R:FR-treated plants when exposed to volatiles emitted by non-infested or *Spodoptera littoralis-*infested low R:FR treated maize plants. Low R:FR-treated emitter plants infested with ten second-instar *S. littoralis* larvae or left uninfested were connected to the glass chambers of receiver plants using Teflon tubing (Fig. 1a). The air flow moved from the emitter’s chamber towards the receiver’s chamber. Volatiles simultaneously emitted by emitter and receiver plants were collected from the glass chamber containing the receiver plant through an air outlet connected to the PTR-TOF-MS system. Because all emitter plants were exposed to low R.FR and volatile fluxes were unidirectional, differences in volatile emissions could only be explained by the light treatment of the receivers (low or high R:FR).

Emission of the green leaf volatiles hexenal, hexen-1-ol, and hexenyl acetate, as well as indole, monoterpenes, sesquiterpenes, and the homoterpenes DMNT and TMTT by *S. littoralis*-infested emitter + intact receiver plants significantly increased over time in comparison to non-infested emitter + intact receiver plants (Fig. 1b). Indole and sesquiterpenes emissions were significantly higher when receiver plants were exposed to low R:FR and volatiles from *S. littoralis*-infested emitter plants than to the same volatiles and high R:FR (Fig. 1b). A similar pattern was observed for monoterpenes and DMNT. As emitter plants were all grown under low R:FR, showed similar levels of herbivore damage (Supplemental Fig. S3), and did not receive any volatile cues from the receiver plants, these differences must be the result of changes in the receivers’ volatile emissions. Thus, FR-enriched light environment enhances maize volatile emissions in response to HIPVs emitted by herbivore-infested neighbors.

### Far-red enrichment boosts maize responses to (Z)-3-hexenyl acetate

Green leaf volatiles such as (*Z*)-3-hexenal, (*Z*)-3-hexen-1-ol, and HAC are HIPVs emitted by maize that can induce the emission of volatiles in neighboring intact maize plants (Farag & Paré, 2002; Engelberth *et al*., 2004). We next tested whether low R:FR specifically modulates maize responses to HAC. We exposed 11-day old B73 maize seedlings to low or high R:FR for two days by modulating FR levels and determined their volatile emissions by using the PTR-TOF-MS system during HAC or clean air exposure and subsequent simulated herbivory (Fig. 2a). These analyses focused on indole and terpenoid emissions because these compounds responded to low R:FR and HIPVs in the previous experiment.

Exposure to HAC induced volatile release in both high and low R:FR-treated maize plants (Fig. 2b). Low R:FR significantly enhanced HAC-induced emissions of monoterpenes and DMNT, while the opposite was observed for indole.

Pre-exposure to HAC increased the emission of all the volatiles signatures after simulated herbivory (priming effect) in both light treatments (Fig. 2b). Emission of DMNT was significantly higher in low R:FR-treated plants than in high R:FR-treated plants pre-exposed to HAC. Irrespective of HAC pre-treatment, herbivory-induced plants exposed to low R:FR emitted more monoterpenes, sesquiterpenes, and DMNT than high R:FR-treated plants (Fig. 2b). An additional experiment comparing volatiles emissions in herbivory-induced and unwounded plants confirmed that low R:FR enhances herbivore-induced emissions of monoterpenes, DMNT, and indole, but it reduced constitutive indole emissions (Supplemental Fig. S4).

In addition, experiments with 30-day-old B73 maize plants showed similar responses to low R:FR and HAC (Supplemental Fig. S5). Importantly, in this experiment, tested variations in horizontal PAR levels in our experimental set up did not correlate with the volatile emissions recorded at the sampling time with the strongest HAC effect (Supplemental Fig. S6), confirming FR specific effects on maize responses to HAC.

The PTR-TOF-MS system only detects molecular mass features, while the structural identity of isomers, e.g., different mono-or sesqui-terpenes, cannot be resolved. To determine the identity of the induced volatile compounds and confirm the observed patterns with an established, orthogonal method, we collected plant volatiles for periods of 2 h for 12 h using Super-Q traps followed by GC-MS analysis of volatile extracts (Supplemental Fig. S7a). HAC exposure significantly increased the emissions of indole, the monoterpene alcohol linalool, the sesquiterpenes (*E*)-β-farnesene and (*E*)-α-bergamotene, and the homoterpene DMNT (Supplemental Fig. S7b). Low R:FR alone had a positive effect on the emission of the two sesquiterpenes and it increased HAC-induced emission of indole, linalool, and (*E*)-α-bergamotene (Supplemental Fig. S7b). We did not detect a significant induction of the monoterpenes α-pinene and β-ocimene. Further PTR-TOF-MS analysis of pure linalool, which is a monoterpene alcohol, showed that the most abundant fragment ion of linalool has a m/z of 137 (C_10_H_17_^+^) (Supplemental Fig. S8) like other monoterpenes. Thus, the induced monoterpene signal detected in the PTR-MS experiments upon HAC and FR light treatment is likely due to enhanced linalool emissions. FR effects on indole emissions agreed with the enhanced induction observed in Fig. 1b but differed from PTR-TOF-MS data displayed in Fig. 2b, indicating a strong context-dependent variation in the emission of this compound. Upon simulated herbivory, linalool, (*E*)-β-farnesene, and (*E*)-α-bergamotene emissions were higher under low R:FR than under high R:FR irrespective of HAC pre-treatment. TMTT was below the limit of detection of the GC-MS. During HAC exposure and after simulated herbivory, DMNT emissions were higher under low R:FR than under high R:FR, albeit these differences were not statistically significant. This might be explained by the lower number of sampling times compared to the PTR-TOF-MS experiments, resulting in a reduced statistical power.

Together, these results demonstrate that FR-enrichment increases maize responsiveness to HAC resulting in higher emission of terpenes in both 11-and 30-day-old maize plants.

### Far-red enrichment enhances JA-Ile accumulation in response to (Z)-3-hexenyl acetate

Volatile emissions in maize can be modulated by jasmonates and ABA (Bruinsma *et al*., 2009; Seidl-Adams *et al*., 2015). To investigate the potential mechanisms by which short-term exposure to low R:FR enhances volatile emissions, levels of these stress-related hormones were analyzed in 11-day old B73 plants exposed to low or high R:FR for two days followed by exposure to HAC or clean air and simulated herbivory (Fig. 3a).

Exposure to HAC increased OPDA, JA, JA-Ile, and ABA levels in both light regimes (Fig. 3b). At 45 min, however, low R:FR-treated plants showed higher HAC-induced JA-Ile levels when compared to high R:FR-treated plants. Low R:FR did not affect constitutive JA, JA-Ile, and SA levels, but it slightly reduced OPDA concentrations and enhanced ABA levels.

Upon simulated herbivory, HAC pre-treated plants displayed higher OPDA and JA-Ile levels when compared to plants pre-exposed to clean air in both light treatments (Fig. 3c). Irrespective of HAC pre-treatment, low R:FR enhanced JA, JA-Ile, and ABA levels at 180 min after simulated herbivory (Fig. 3c).

### Far-red enrichment modulates the expression of volatile biosynthetic genes

Next, we tested whether low R:FR-induced emission of volatiles in B73 seedlings correlated with changes in the expression of terpene-related biosynthetic genes during HAC exposure and subsequent simulated herbivory (Fig. 4a).

Exposure to HAC significantly induced the expression of *FPSS3,* which provides precursors for sesquiterpene biosynthesis (Richter *et al*., 2015), *TPS10*, involved in the biosynthesis (*E*)-α-bergamotene and (*E*)-β-farnesene (Schnee *et al*., 2006), and *TPS2*, involved in the production of linalool, DMNT, and TMTT (Richter *et al*., 2016) (Fig. 4b). *TPS1*, involved in the biosynthesis of (*E*)-β-farnesene, linalool derivatives, and *ent-*kaurene, the latter serving as an intermediate in the production of kauralexins (Schnee *et al*., 2002; Fu *et al*., 2016; Xu *et al*., 2019), was also induced by HAC. Expression of *CYP92C5*, involved in the production of DMNT (Richter *et al*., 2016), was slightly reduced by HAC at 180 min after the start of the treatment (Fig. 4b). HAC effects were not influenced by light. However, when compared to high R:FR treated plants, low R:FR-treated plants showed higher *TPS1* expression levels at 45 and 90 min, and reduced expression of *CYP92C5* and *FPPS3* genes at 45 min and 90 min, respectively, after the start of the experiment (Fig. 4b).

Upon simulated herbivory, HAC pre-treatment enhanced the expression of *TPS2* and *TPS10* and suppressed *CYP92C5* expression in both low and high R:FR-treated plants, with no effect on *FPSS3* and *TPS1* (Fig. 4c). Low R:FR led to a stronger HAC-mediated *CYP92C5* suppression and induction of *TPS10* at 45 and 90 min, respectively, upon simulated herbivory. Irrespective of HAC pre-treatment, low R:FR slightly enhanced *TPS2*, *TPS10*, and *CYP92C5* expressions after simulated herbivory (Fig. 4c).

These results show that FR-enriched environment can modulate the expression of some, but not all, terpene biosynthesis genes, and that these slight changes are unlikely to fully explain the differences in volatile release in response to HAC.

### Far-red enrichment increases stomatal conductance and photosynthesis

Emission of sesquiterpenes can be regulated by light-dependent stomata movements in maize (Seidl-Adams *et al*., 2015). We next tested whether differences in stomata conductance accounted for the increase in volatile emissions under low R:FR. For this, we exposed 11-day-old B73 seedlings to low or high R:FR for two days and determined leaf stomatal conductance in two maize leaves per plant at 3 h after the start of HAC exposure and at 4 h after subsequent simulated herbivory (Fig. 5a). To differentiate the effects of wounding on stomatal conductance, we included unwounded plants in this experiment.

**Figure 5.**
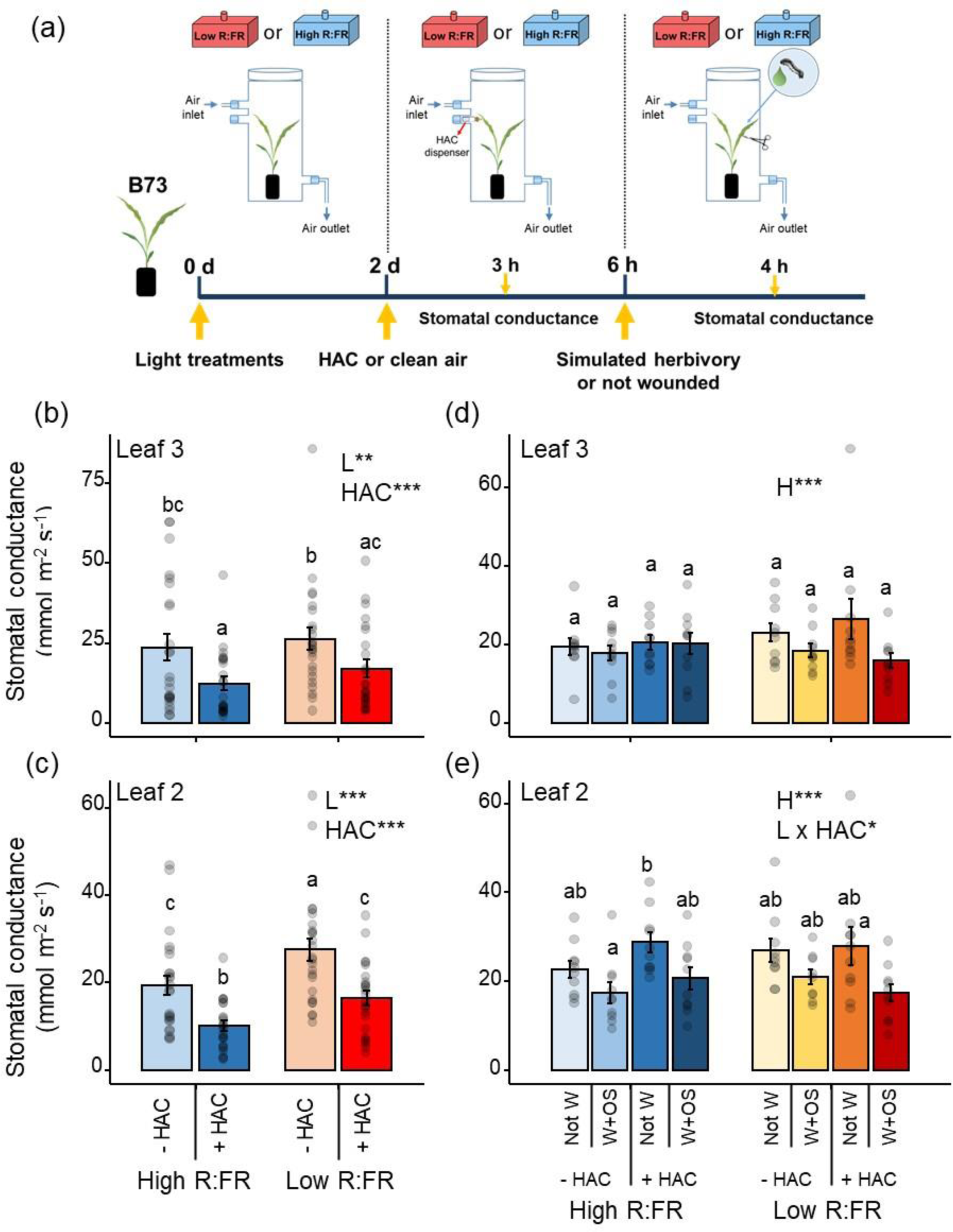
Far-red enrichment increases stomatal conductance in maize. **(a)** Schematical overview of the experimental set up and sampling times. Stomatal conductance (mean ± SEM, *n* = 20) determined in the abaxial side of **(b)** leaf #3 and **(c)** leaf #2 from the bottom of low and high R:FR-treated B73 maize (*Zea* mays) plants after 3 h of exposure to (*Z*)-3-hexenyl acetate (HAC) or clean air. Effects of light treatment (L), HAC exposure and their interaction were tested by two-way ANOVAs. Stomatal conductance (mean ± SEM, *n* = 10) determined in the abaxial side of **(d)** leaf #3 and **(e)** leaf #2 of low and high R:FR-treated B73 plants that were induced with simulated herbivory (wounding and application of *Spodoptera littoralis* oral secretions; W+OS) or left unwounded (Not W) after exposure to HAC or clean air for 6 h. Measurements were performed close to the wounding sites at 4 h after simulated herbivory. Effects of light treatment (L), HAC exposure, herbivory (H), and their interaction were tested by three-way ANOVAs. In all graphs, different letters denote significant differences among groups tested by Tukey-HSD post hoc test. Statistically significant effects are shown in each graph (\**p* < 0.05, ** *p* < 0.001, *** *p* < 0.001).

Exposure to HAC significantly reduced stomatal conductance in maize (Fig. 5bc). Irrespective of HAC, low R:FR significantly increased stomatal conductance, which was not explained by changes in stomatal densities (Supplemental Fig. S9). We did not detect significant interactions between light and HAC on stomatal conductance.

Simulated herbivory reduced stomatal conductance close to the wounding sites when compared to unwounded controls (Fig. 5de), a phenomenon reported to be driven by OPDA signaling in local leaves of dicotyledonous plants (Savchenko et al., 2014; Meza-Canales et al., 2017). In leaf 2, stomatal closure was slightly stronger in wounded plants exposed to low R:FR and HAC pre-treatment (Fig. 5e).

Stomatal opening as well as volatile biosynthesis are tightly coupled to photosynthetic rates (Arimura *et al*., 2008; Way & Pearcy, 2012). We therefore evaluated if light and volatile cues interact to regulate photosynthesis (Fig. 6). While HAC had no significant effect on photosynthesis, photosynthetic rates were significantly increased under low R:FR in leaf 3 (Fig. 6b).

**Figure 6.**
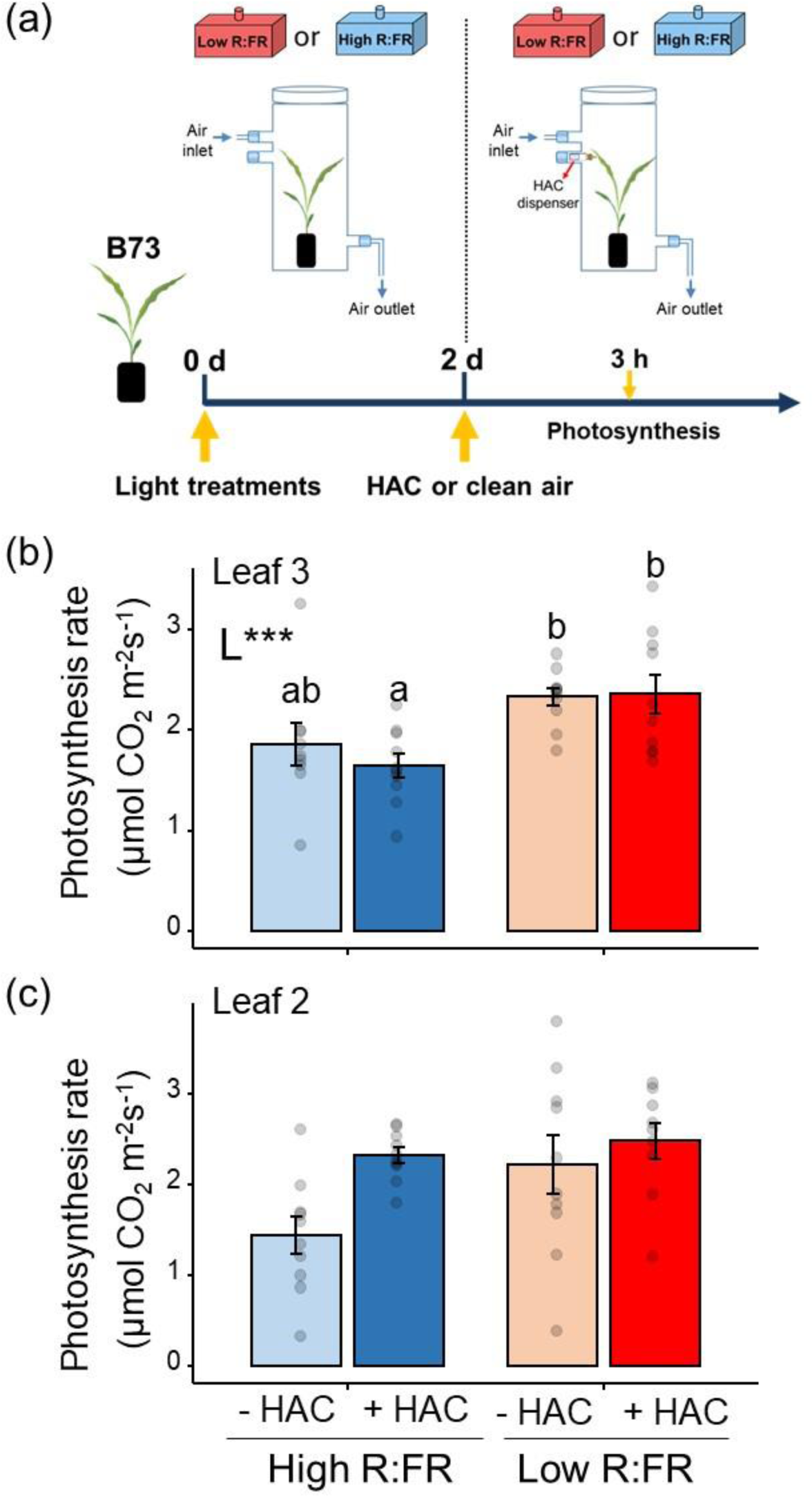
Far-red enrichment increases photosynthesis in maize. **(a)** Schematical overview of the experimental set up and sampling times. Photosynthesis rate (mean ± SEM, *n* = 10) determined in the abaxial side of **(a)** leaf 3 and **(b)** leaf 2 from the bottom of high and low R:FR-treated B73 maize plants at 3 h after the start of exposure to (*Z*)-3-hexenyl acetate (HAC) or clean air. Effects of light treatment (L) and HAC on stomatal density were tested by two-way ANOVAs. Statistically significant effects are shown in the graphs. \*\*\**p* < 0.001. Different letters denote significant differences among groups tested by Tukey-HSD post hoc test at *p* < 0.05.

### Maize volatile emissions in response to (Z)-3-hexenyl acetate are reduced in the phyB1phyB2 mutant

Low R:FR is sensed by the phytochrome family of photoreceptors, of which phyB is the central regulator in Arabidopsis (Ballaré & Pierik, 2017). In maize, gene duplication has resulted in two PhyB1 and PhyB2 genes, both involved in FR signaling (Sheehan *et al*., 2007; Dubois *et al*., 2010). We next determined whether a non-functional phyB enhances maize volatile emissions in respoinse to HAC like low R:FR does.

First, we exposed 11-day-old *phyB1phyB2* maize double mutant plants and its wild-type (inbred line W22) to high R:FR for two days and determined volatile emissions during HAC exposure and after simulated herbivory using the PTR-TOF-MS system (Fig. 7a). Compared to the wild-type, constitutive emission of indole, monoterpenes, sesquiterpenes, and DMNT was significantly lower in *phyB1phyB2* plants (Fig. 7b). HAC increased volatile emissions in wild-type and *phyB1phyB2* plants, but the induction of volatiles was significantly lower in *phyB1phyB2* plants compared to the wild-type. Similar patterns were observed following simulated herbivory, in the priming phase.

**Figure 7.**
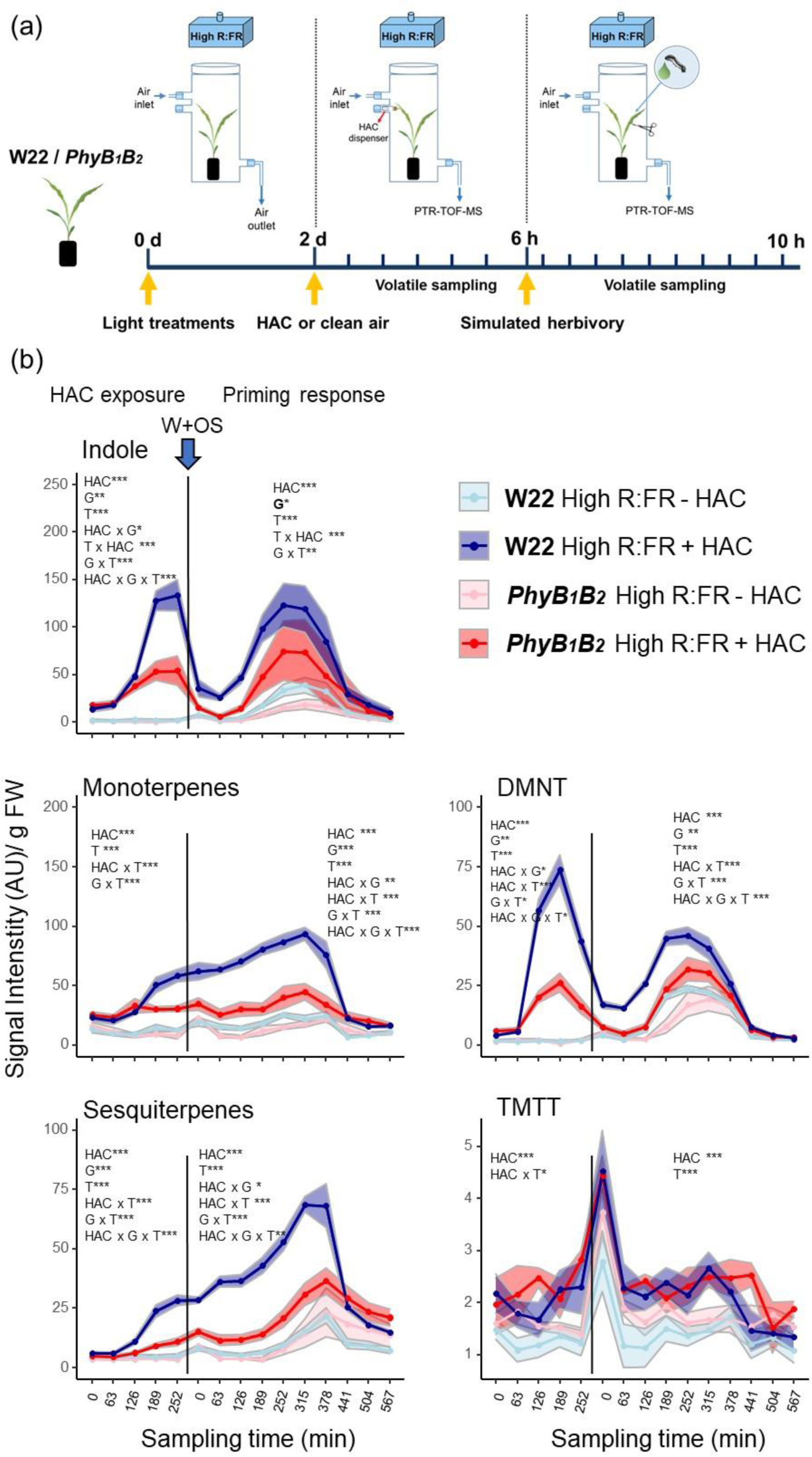
Maize volatile emissions in response to (*Z*)-3-hexenyl acetate are reduced in the *phyB1phyB2* mutant. **(a)** Schematical overview of the experimental set up and sampling times **(b)**Time-series analysis of volatile emissions (mean ± SEM, *n* = 6) determined by PTR-TOF-MS in high R:FR-treated *PhyB_1_B_2_* double mutant and wild-type (inbred line ‘W22’) maize (*Zea mays*) plants during exposure to (*Z*)-3-hexenyl acetate (+ HAC) or clean air (-HAC) and after induction with simulated herbivory (wounding and application of *Spodoptera littoralis* oral secretions; W+OS) (priming phase). HAC exposure and volatile measurements started between 9:00 and 10:00 am. The effects of the sampling time (T), HAC, genotype (G), and their interactions on volatile emissions were tested separately for the HAC exposure and priming phase using linear mixed effects models. Statistically significant effects are shown in each graph (\**p* < 0.05, ** *p* < 0.001, *** *p* < 0.001). DMNT and TMTT stand for the homoterpenes (*E*)-4,8-dimethyl-1,3,7-nonatriene and (*E*, *E*)-4,8,12-trimethyltrideca-1,3,7,11-tetraene, respectively. AU refers to arbitrary units.

To better understand the unexpected response of *phyB1phyB2* plants, we conducted additional experiments. First, we analyzed photomorphogenic responses of *phyB1phyB2* and wild-type W22 plants to low R:FR. As expected, *phyB1phyB2* plants grew longer stem internodes irrespective of the light regime (Supplemental Fig. S10). The wild-type W22 did not show a strong response to low R:FR, which might be explained by maize genotype-dependent responses to supplemental FR (Dubois *et al*., 2010).

Next, we compared the responsiveness of the wild-type W22 and *phyB1phyB2* plants to HAC followed by simulated herbivory when grown under high or low R:FR. Low R:FR significantly enhanced HAC-induced emission of monoterpenes, sesquiterpenes, and DMNT in wild-type W22 plants (Supplemental Fig. S11) as observed before in B73 (Fig. 2b and Supplemental Fig. S7), whereas the effect on indole differed along the time course. Light did not significantly affect volatile emissions upon simulated herbivory nor priming responses in W22. In *phyB1phyB2* plants, low R:FR slightly increased indole, monoterpenes, and sesquiterpenes emissions at the start of the HAC treatment (Supplemental Fig. S12), and it did not significantly affect volatile emissions after simulated herbivory.

These experiments indicate that *phyB1phyB2* plants retain partial, but not full, enhanced responsiveness to HAC when simultaneously exposed to low R:FR.

### Maize volatile responsiveness to (Z)-3-hexenyl acetate is stronger under short-term than under long-term exposure to far-red enrichment

Why do *phyB1phyB2* mutant plants that behave as if continuously exposed to low R:FR show suppression of HAC-induced volatile emissions? One possible explanation is that phyB is inactivated in *phyB1phyB2* mutant plants over the full course of their development, while in FR experiments plants have a deactivated phytochrome for only two days preceding volatile exposure. To test whether duration of phyB inactivation matters for volatile responsiveness, seven-day-old W22 plants were exposed to low or high R:FR for 6 days, or 4 days of high R:FR followed by 2 days of low R:FR (Fig. 8a). Then, we monitored volatile emissions during HAC exposure and subsequent simulated herbivory using the PTR-TOF-MS system (Fig. 8b).

**Figure 8.**
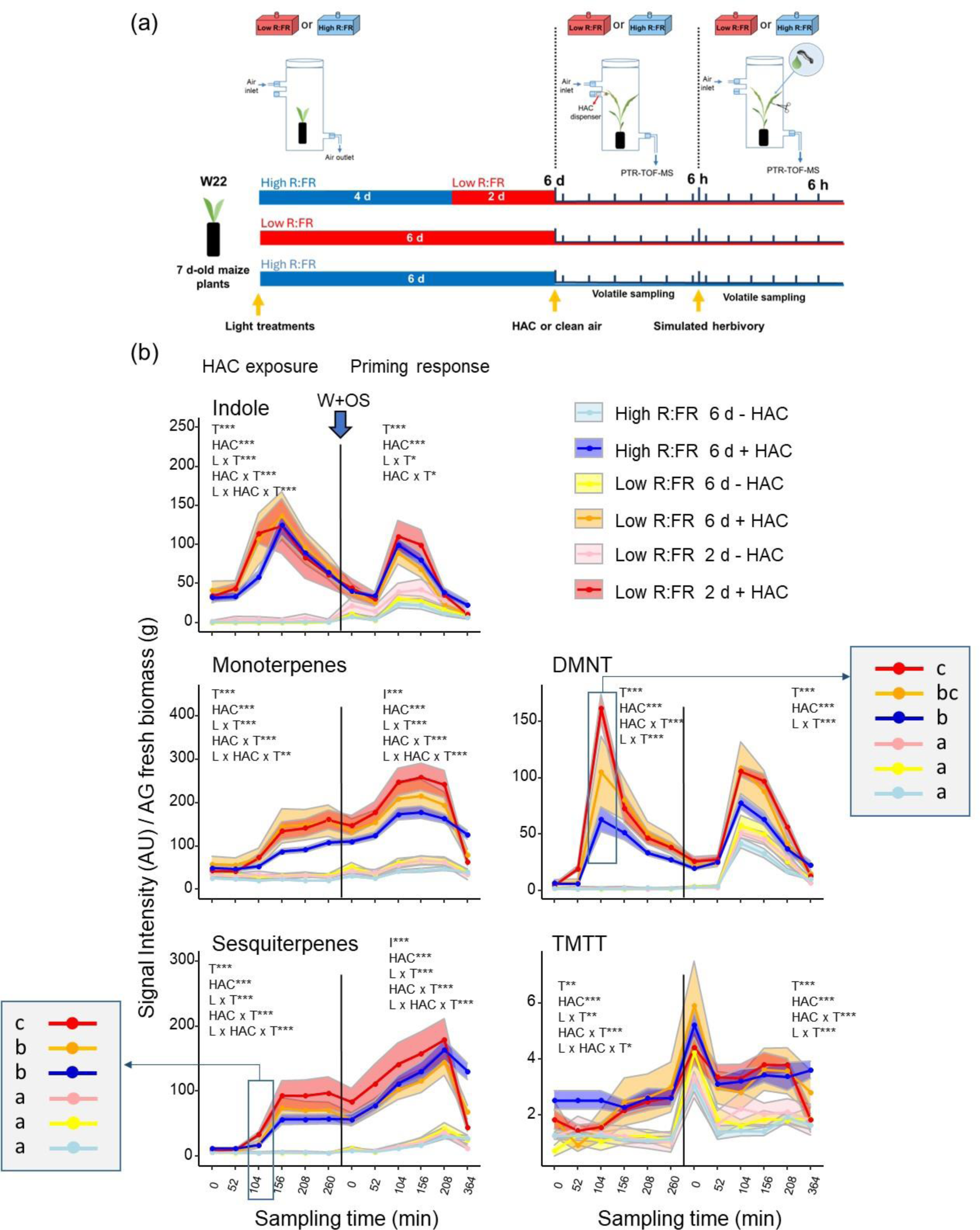
Maize volatile responsiveness to (*Z*)-3-hexenyl acetate is stronger under short-than under long-term far-red enrichment. **(a)** Schematic overview of the experimental set-up. Seven-day-old W22 maize (*Zea mays*) seedlings were exposed to: (1) high R:FR for 6 days, (2) low R:FR for 6 days, or (3) high R:FR for 4 days and transferred to low R:FR conditions for 2 days. Thereafter, plants were exposed to (*Z*)-3-hexenyl acetate (HAC) or clean air for 6 h and subsequently all induced with simulated herbivory (wounding and application of *Spodoptera littoralis* oral secretions; W+OS). **(b)** Time-series analysis of volatile emissions (mean ± SEM, *n* = 6) determined by PTR-TOF-MS during exposure to HAC (+ HAC) or clean air (-HAC) and after induction with simulated herbivory (priming phase). HAC exposure and volatile measurements started between 9:00 and 10:00 am. The effects of the sampling time (T), HAC, light treatment (L) and their interactions on volatile emissions were tested separately for the HAC exposure and priming phase using linear mixed effects models. Statistically significant effects are shown in each graph (\**p* < 0.05, ** *p* < 0.001, *** *p* < 0.001). Shaded grey squares containing different letters indicate significant differences among treatments tested by estimated marginal means at *p* < 0.05. DMNT and TMTT stand for the homoterpenes (*E*)-4,8-dimethyl-1,3,7-nonatriene and (*E*, *E*)-4,8,12-trimethyltrideca-1,3,7,11-tetraene, respectively. AU refers to arbitrary units.

Wild-type W22 plants exposed to low R:FR for 2 days showed stronger volatile responses to HAC than plants exposed to high R:FR (Fig. 8b) as in previous experiments (Supplemental Fig. S11). Short or long-term exposure to low R:FR induced higher indole, sesquiterpenes, and DMNT emissions, but the inverse pattern was observed for TMTT. Induced sesquiterpenes and DMNT emissions at early times of HAC exposure were lower in plants exposed to low R:FR for 6 days than to shorter low R:FR exposure (2 days) and high R:FR only (Fig. 8b).

Upon herbivory, short-term exposure (2 days) to low R:FR also enhanced monoterpenes and sesquiterpenes emissions, but this response was attenuated under long-term low R:FR exposure (6 days). Thus, long-term inactivation of phyB by low R:FR may partially explain the discrepancy between the *phyB1phyB2* double mutant phenotype and short-term FR supplementation experiments. Hence, these results suggest that short-and long-term inactivation of phyB might result in different physiological responses.

## Discussion

In closing or dense canopies, plants can detect the proximity and physiological state of neighboring plants through light and volatile cues. Our results demonstrate that maize plants integrate both stimuli into distinct volatile signaling patterns. Below, we discuss the potential mechanisms and ecological consequences of this phenomenon.

Herbivore-induced plant volatiles can induce defense-related responses in undamaged plants including the biosynthesis and emission of volatiles (Arimura *et al*., 2002; Choh *et al*., 2004; Engelberth *et al*., 2004) and the priming of defense responses when actual herbivory occurs (Ton *et al*., 2007; Frost *et al*., 2008; Erb *et al*., 2015; Hu *et al*., 2019; Paudel Timilsena *et al*., 2020). Here, we report that a FR-enriched light environment prompts maize plants to respond more strongly to HIPVs emitted by their neighbors. We found that volatile release in response to HIPVs or HAC is enhanced in plants exposed to FR-enriched conditions, the typical spectral signature of the light environment in closing plant canopies and shaded environments. In line with our findings, Chautá and Kessler (2022) also reported a higher emission of a sesquiterpene in tall goldenrod (*Solidago altissima*) plants exposed to supplemental FR and HIPVs from neighbors, and FR alone has been shown to increase the emissions of some VOCs in tomato (Cortés *et al*., 2016), albeit the mechanisms remained to be determined.

Most of the volatile identities and PTR-TOF-MS emission patterns in the present work were validated using a complementary dynamic volatile collection and GC-MS detection. The higher sensitivity of the PTR-TOF-MS allowed the detection of less abundant volatiles such as TMTT, and it also enhanced the statistical power of the analyses by increasing their temporal resolution. Strikingly, HAC-induced indole emissions showed varying patterns between experiments, being slightly enhanced by FR in some of them during the time-series analyses while strongly reduced in others (Fig. 2b). Further experimentation is needed to determine the context-dependent regulation of indole emissions.

Plant responses to volatiles are regulated by the jasmonate signaling pathway (Arimura *et al*., 2000; Hu, 2022) as part of an integrated signaling network that controls plant growth and defense responses (Schmelz *et al*., 2003; Bosch *et al*., 2014; Mujiono *et al*., 2021). In maize, HAC induces the accumulation of jasmonates and expression of JA-related marker genes before and after herbivory induction (Hu *et al*., 2019). Here, we confirmed that HAC strongly up-regulated JA signaling in maize. Furthermore, we found that a FR-enriched environment resulted in a two-fold higher induction of JA-Ile levels during HAC exposure. This is surprising considering the reported suppressing effects of low R:FR on JA accumulation (Fernandez-Milmanda et al., 2020) and signaling in Arabidopsis (Cerrudo *et al*., 2012; de Wit *et al*., 2013; Chico *et al*., 2014; Leone *et al*., 2014). Yet, low R:FR induced characteristic shade-associated morphological responses in maize, such as the elongation of stems. Furthermore, we observed strong inductions of auxin and ABA levels in low R:FR treated plants, with both hormones involved in shade-induced suppression of branching (tillering in grasses) (Küpers *et al*., 2020; Fernández-Milmanda & Ballaré, 2021). PAR levels and R:FR ratios used in our work were similar to conditions employed in previous work reporting the effects of supplemental FR on hormonal signaling and defenses in Arabidopsis (Pierik *et al*., 2009; Wit *et al*., 2013) and tomato (Courbier *et al*., 2020; Courbier *et al*., 2021). Thus, although maize responses to shade show similarities with those described for Arabidopsis and other plant species, there might be significant points of divergence. Additional analyses using reverse genetics and pharmacological approaches can reveal the importance of the enhanced jasmonate response and the potential role of auxin and ABA in maize responses to HAC.

Plant volatile emission is controlled by biochemical, physiological, and physicochemical processes including the rate of volatile biosynthesis, the activities of enzymes, the volatility and diffusion rates of distinct volatile compounds, and stomatal conductance (Niinemets *et al*., 2004; Seidl-Adams *et al*., 2015). Our study showed that low R:FR changed the expression of a subset of volatile biosynthesis genes in response to HAC and simulated herbivory. However, the observed interactions did not fully track the changes in volatile emissions during HAC exposure. Thus, it is unlikely that light and volatile cues interact only at this level to boost volatile release in maize. Low R:FR effects on the diffusion of volatiles through the cuticle is also an unlikely explanation, as the cuticular diffusion coefficients of monoterpenes is lower than the diffusion coefficient of air and water, indicating that the cuticle provides an efficient barrier for these plant volatiles (Niinemets, 2003). We therefore reasoned that volatile emissions might be additionally modulated by induced changes in stomatal conductance. Our results showed that both exposure to HAC and simulated herbivory reduced leaf stomatal conductance, which contrasted with the strong increase in volatile emissions. This might be explained by the fact that leaf stomata can only limit volatile emission if the decrease in stomatal conductance (i.e., closure) is larger than the increase in the volatile diffusion gradient (Lin *et al*., 2022). Most probably, HAC induction of volatile biosynthesis quickly increased their concentration in the intercellular air spaces, enhancing volatile diffusion gradients and overruling stomatal limitations.

Interestingly, FR-enriched environment had a positive effect on leaf stomatal conductance. This is consistent with previous findings in bean (*Phaseolus vulgaris*) where supplemental FR added from above to sub-saturating levels of visible light increased stomatal conductance (Holmes *et al*., 1986). More recently, FR has been shown to enhance photochemical efficiency and photosynthesis in several dicotyledonous and C3 and C4 monocotyledonous species, including maize, when added to a background of white light (Zhen & Bugbee, 2020). In maize, FR photons were recently shown to account for a large fraction of the photosynthetic activity under vegetation shade (Zhen *et al*., 2022). This was explained by their enhancing linear electron transport efficiency and cyclic electron flow around the photosystem I (Zhen *et al*., 2022). In C4 species such as maize, this is especially relevant due to the extra ATP requirement for the CO_2_-concentrating mechanism in the bundle sheath cells (Johnson, 2011). Hence, our results align with the expected stomatal and photosynthetic responses of maize plants in dense canopies characterized by low irradiances in the visible light range (400-700 nm) and enrichment in the FR region of the spectrum. Zhen and Bugbee (2020) also found that the addition of equal amounts of FR or white photons causes similar increases in net photosynthesis, while the increase in stomatal conductance was comparatively smaller under FR. This suggested that stomatal responses to FR were most likely a response to the increase in photosynthetic rate. Stomatal conductance in Arabidopsis *phyB* mutants have been shown to be modulated by changes in ABA sensitivity under drought stress (González *et al*., 2012). Whether the increase in stomatal conductance from adding FR result from increased photosynthetic rates and/or of alteration of ABA sensitivity needs to be established. Yet, it is conceivable that the effects of FR-enriched environment on the emission of volatiles in response to HAC might be explained, at least in part, by the positive effects of supplemental FR on stomatal conductance and/or higher photosynthetic rates. The latter might affect the availability of precursors for volatile biosynthesis (Arimura *et al*., 2008).

Unlike FR supplementation experiments, *phyB1phyB2* plants displayed a diminished response to HAC. These results align with the suppressive effect of phyB inactivation on plant defenses (Ballaré, 2014; Pierik & Ballaré, 2021). We reasoned that long-term phyB inactivation (as present in the *phyB1phyB2* mutant) may have different effects from those triggered by a short-term inactivation via changes in R:FR. Indeed, longer exposure of maize seedlings to low R:FR resulted in a slightly weaker effect of low R:FR on HAC-induced volatile emissions. At the same time, the *phyB1phyB2* mutant retained some ability to respond to low R:FR with higher HAC induced volatile emissions, reinforcing the potential effects of FR on the photosynthetic machinery and stomatal conductance. We therefore conclude that (i) the duration of phyB inactivation might explain FR effects on maize responses to HAC, and that (ii) an enhanced photosynthetic efficiency and stomatal conductance, and/or (iii) other FR photoreceptors may also play a role in this phenomenon. Under deep shade (R:FR < 0.8), for instance, phyA is strongly activated and prevents the excessive elongation of seedlings in Arabidopsis, antagonizing phyB FR-induced responses (Casal *et al*., 2014; Martínez-García *et al*., 2014; Lim *et al*., 2018; Yang *et al*., 2018). Furthermore, PhyA promotes photosynthetic processes in rice (Panda *et al*., 2023). PhyA therefore represents a candidate that may be involved in the responses to light and volatile cues in maize.

From a biological perspective, the capacity to integrate light and volatile cues would seem particularly beneficial to fend off an attacking herbivore. While induced volatiles can serve as reliable cues for the presence of herbivores on neighboring plants (Moreira *et al*., 2018; Meents & Mithöfer, 2020), light cues can provide information about the proximity of neighbors and, therefore, the risk of being attacked next. Because the effects of herbivore attack are more severe under competition (Vries *et al*., 2018), it might be advantageous for a plant to respond more strongly to induced volatiles in the presence of neighboring plants and vegetative shade. The enhanced emission of volatiles might, for instance, promote the attraction of natural enemies in a setting where their recruitment is a competitive struggle (Kessler & Baldwin, 2001; Cortés et al., 2016; Aartsma *et al*., 2017). Plant volatiles may also decrease ovipositing of gravid female moths (Moraes *et al*., 2001). However, these volatiles may also attract herbivores themselves (Mérey *et al*., 2013), in which case becoming “apparent” in a dense stand would be the wrong strategy. Further studies on the adaptive benefits of volatile responses in dense stands will help us to understand the evolutionary and adaptive character of plant responses to light and volatile cues in agricultural and ecological settings. Clearly, the integration of light and volatile cues deserves attention as a potentially major determinant of plant-plant, plant-herbivore, tri-trophic interaction dynamics, and community structure.

## Acknowledgments

We thank Prof. Dr. Tobias Züst for his help in the optimization of GC-MS protocols and the identification of maize volatile compounds. We also thank Prof. Dr. Christian Fankhauser and Dr. Martina Legris for their help with light spectral measurements. The work of BCJS was supported by the Marie Sklodowska-Curie Action Individual Fellowship (European Union Horizon 2020, Grant Nr. 794,947). The work of LW was supported by the Marie Sklodowska-Curie Action Individual Fellowship (European Union Horizon 2020, Grant Nr. 886651). This work was supported by the Velux Foundation (Grant Nr. 1231), the Swiss National Science Foundation (Grant. Nr. 200355), the State Secretariat for Education, Research, and Innovation SERI (Project CANWAS) and the University of Bern.

## Author contributions

ME conceived the study. ME, RE-B, BCJS, CB, and CAMR designed experiments. RE-B conducted the experiments. RE-B performed the PTR-TOF-MS measurements and analyses. LW assisted with PTR-TOF-MS measurements and conducted the analysis of linalool emissions from dispensers. RE-B and BCJS conducted gene expression analyses. RE-B and GG performed hormone analyses. RE-B and YZ conducted stomatal measurements. RE-B conducted photosynthesis measurements. RE-B analyzed the data. RE-B, BCJS, CB and ME interpreted the data. RE-B and ME wrote the manuscript and all the authors contributed to revisions.

## Data availability

The data generated for this manuscript will be archived in Dryad and the data DOI will be included at the end of the article.

## Supporting Information

**Figure S1.** Light spectral composition determined in far-red supplementation experiments.

**Figure S2.** Effects of far-red enrichment on maize growth and auxin levels.

**Figure S3.** Aboveground biomass in maize plants infested with *Spodoptera littoralis* larvae.

**Figure S4.** Effect of far-red enrichment on herbivory-induced plant volatile emission in maize

**Figure S5.** Effect of far-red enrichment and (*Z*)-3-hexenyl acetate on volatile emissions in 30-day-old maize plants.

**Figure S6** Spatial variations in photosynthetically active radiation (PAR) levels in the experimental set up did not influence 30-day-old maize responses to (*Z*)-3-hexenyl acetate.

**Figure S7.** Effect of far-red enrichment on maize volatile emission during exposure to (*Z*)-3-hexenyl acetate and after simulated herbivory measured via GC-MS methods.

**Figure S8.** Measurement of pure linalool in the PTR-TOF-MS system.

**Figure S9.** Effect of far-red enrichment and (*Z*)-3-hexenyl acetate on stomatal density.

**Figure S10.** Effect of far-red enrichment on the growth of *phyB1phyB2* maize double mutant and its wild-type.

**Figure S11.** Effect of far-red enrichment on volatiles emission during exposure to (*Z*)-3-hexenyl acetate and after simulated herbivory in the maize inbred line W22.

**Figure S12.** Effect of far-red enrichment on volatiles emission during exposure to (*Z*)-3-hexenyl acetate and after simulated herbivory in the *phyB1phyB2* maize double mutant.

**Supplementary Method S1.** Detailed protocol for phytohormone extraction and analysis.

**Supplementary Table S1.** Primer’s specifications for gene expression analyses

## References

Aartsma Y, Bianchi FJJA, van der Werf W, Poelman EH, Dicke M. 2017. Herbivore-induced plant volatiles and tritrophic interactions across spatial scales. New Phytologist 216: 1054–1063.

Alba JM, Schimmel BCJ, Glas JJ, Ataide LMS, Pappas ML, Villarroel CA, Schuurink RC, Sabelis MW, Kant MR. 2015. Spider mites suppress tomato defenses downstream of jasmonate and salicylate independently of hormonal crosstalk. The New phytologist 205: 828–840.

Arimura G, Köpke S, Kunert M, Volpe V, David A, Brand P, Dabrowska P, Maffei ME, Boland W. 2008. Effects of feeding Spodoptera littoralis on lima bean leaves: IV. Diurnal and nocturnal damage differentially initiate plant volatile emission. Plant Physiology 146: 965–973.

Arimura G, Ozawa R, Nishioka T, Boland W, Koch T, Kühnemann F, Takabayashi J. 2002. Herbivore-induced volatiles induce the emission of ethylene in neighboring lima bean plants. The Plant journal: for cell and molecular biology 29: 87–98.

Arimura G, Ozawa R, Shimoda T, Nishioka T, Boland W, Takabayashi J. 2000. Herbivory-induced volatiles elicit defence genes in lima bean leaves. Nature 406: 512–515.

Ballaré CL. 2009. Illuminated behaviour: phytochrome as a key regulator of light foraging and plant anti-herbivore defence. Plant, Cell & Environment 32: 713–725.

Ballaré CL. 2014. Light regulation of plant defense. Annual review of plant biology 65: 335–363.

Ballaré CL, Pierik R. 2017. The shade-avoidance syndrome: multiple signals and ecological consequences. Plant, Cell & Environment 40: 2530–2543.

Ballaré CL, Scopel AL, Sánchez RA. 1990. Far-red radiation reflected from adjacent leaves: an early signal of competition in plant canopies. Science 247: 329–332.

Barbosa P, Hines J, Kaplan I, Martinson H, Szczepaniec A, Szendrei Z. 2009. Associational Resistance and Associational Susceptibility: Having Right or Wrong Neighbors. Annual Review of Ecology, Evolution, and Systematics 40: 1–20.

Block AK, Hunter CT, Rering C, Christensen SA, Meagher RL. 2018. Contrasting insect attraction and herbivore-induced plant volatile production in maize. Planta 248: 105–116.

Bosch M, Wright LP, Gershenzon J, Wasternack C, Hause B, Schaller A, Stintzi A. 2014. Jasmonic acid and its precursor 12-oxophytodienoic acid control different aspects of constitutive and induced herbivore defenses in tomato. Plant Physiology 166: 396–410.

Bruinsma M, Posthumus MA, Mumm R, Mueller MJ, van Loon JJA, Dicke M. 2009. Jasmonic acid-induced volatiles of Brassica oleracea attract parasitoids: effects of time and dose, and comparison with induction by herbivores. Journal of Experimental Botany 60: 2575–2587.

Campos ML, Yoshida Y, Major IT, Oliveira Ferreira D de, Weraduwage SM, Froehlich JE, Johnson BF, Kramer DM, Jander G, Sharkey TD et al. 2016. Rewiring of jasmonate and phytochrome B signalling uncouples plant growth-defense tradeoffs. Nature Communications 7: 12570.

Cargnel MD, Demkura PV, Ballaré CL. 2014. Linking phytochrome to plant immunity: low red: far-red ratios increase Arabidopsis susceptibility to Botrytis cinerea by reducing the biosynthesis of indolic glucosinolates and camalexin. New Phytologist 204: 342–354.

Casal JJ, Candia AN, Sellaro R. 2014. Light perception and signalling by phytochrome A. Journal of Experimental Botany 65: 2835–2845.

Cerrudo I, Keller MM, Cargnel MD, Demkura PV, de Wit M, Patitucci MS, Pierik R, Pieterse CMJ, Ballaré CL. 2012. Low red/far-red ratios reduce Arabidopsis resistance to Botrytis cinerea and jasmonate responses via a COI1-JAZ10-dependent, salicylic acid-independent mechanism. Plant physiology 158: 2042–2052.

Chautá A, Kessler A. Metabolic integration of spectral and chemical cues mediating plant responses to competitors and herbivores. Plants 11(20), 2768.

Chico J-M, Fernández-Barbero G, Chini A, Fernández-Calvo P, Díez-Díaz M, Solano R. 2014. Repression of Jasmonate-Dependent Defenses by Shade Involves Differential Regulation of Protein Stability of MYC Transcription Factors and Their JAZ Repressors in Arabidopsis. The Plant Cell 26: 1967–1980.

Choh Y, Shimoda T, Ozawa R, Dicke M, Takabayashi J. 2004. Exposure of lima bean leaves to volatiles from herbivore-induced conspecific plants results in emission of carnivore attractants: active or passive process? Journal of Chemical Ecology 30: 1305–1317.

Chrétien LTS, David A, Daikou E, Boland W, Gershenzon J, Giron D, Dicke M, Lucas-Barbosa D. 2018. Caterpillars induce jasmonates in flowers and alter plant responses to a second attacker. New Phytologist 217: 1279–1291.

Cortés LE, Weldegergis BT, Boccalandro HE, Dicke M, Ballaré CL. 2016. Trading direct for indirect defense? Phytochrome B inactivation in tomato attenuates direct anti-herbivore defenses whilst enhancing volatile-mediated attraction of predators. New Phytologist 212: 1057–1071.

Courbier S, Grevink S, Sluijs E, Bonhomme P-O, Kajala K, van Wees SCM, Pierik R. 2020. Far-red light promotes Botrytis cinerea disease development in tomato leaves via jasmonate-dependent modulation of soluble sugars. *Plant*, Cell & Environment 43: 2769–2781.

Courbier S, Snoek BL, Kajala K, Li L, van Wees SCM, Pierik R. 2021. Mechanisms of far-red light-mediated dampening of defense against Botrytis cinerea in tomato leaves. Plant Physiology 187: 1250–1266.

Dubois PG, Brutnell TP. 2009. Light Signal Transduction Networks in Maize. In: Bennetzen JL, Hake SC, eds. Handbook of maize. Its biology. New York: Springer, 205–227.

Dubois PG, Olsefski GT, Flint-Garcia S, Setter TL, Hoekenga OA, Brutnell TP. 2010. Physiological and genetic characterization of end-of-day far-red light response in maize seedlings. Plant physiology 154: 173–186.

Engelberth J, Alborn HT, Schmelz EA, Tumlinson JH. 2004. Airborne signals prime plants against insect herbivore attack. Proceedings of the National Academy of Sciences of the United States of America 101: 1781–1785.

Erb M, Flors V, Karlen D, Lange E de, Planchamp C, D’Alessandro M, Turlings TCJ, Ton J. 2009. Signal signature of aboveground-induced resistance upon belowground herbivory in maize. The Plant Journal 59: 292–302.

Erb M, Veyrat N, Robert CAM, Xu H, Frey M, Ton J, Turlings TCJ. 2015. Indole is an essential herbivore-induced volatile priming signal in maize. Nature Communications 6: 6273.

Escobar-Bravo R, Klinkhamer PGL, Leiss KA. 2017. Interactive Effects of UV-B Light with Abiotic Factors on Plant Growth and Chemistry, and Their Consequences for Defense against Arthropod Herbivores. Frontiers in Plant Science 8: 278.

Fankhauser C. 2001. The phytochromes, a family of red/far-red absorbing photoreceptors. Journal of Biological Chemistry 276: 11453–11456.

Farag MA, Paré PW. 2002. C6-Green leaf volatiles trigger local and systemic VOC emissions in tomato. Phytochemistry 61: 545–554.

Fernández-Milmanda GL, Ballaré CL. 2021. Shade Avoidance: Expanding the Color and Hormone Palette. Trends in plant science 26: 509–523.

Fernández-Milmanda GL, Crocco CD, Reichelt M, Mazza CA, Köllner TG, Zhang T, Cargnel MD, Lichy MZ, Fiorucci A-S, Fankhauser C et al. 2020. A light-dependent molecular link between competition cues and defence responses in plants. Nature plants 6: 223–230.

Finch-Savage WE, Leubner-Metzger G. 2006. Seed dormancy and the control of germination. The New phytologist 171: 501–523.

Fiorucci A-S, Fankhauser C. 2017. Plant Strategies for Enhancing Access to Sunlight. Current biology: CB 27: R931–R940.

Franklin KA. 2008. Shade avoidance. The New phytologist 179: 930–944.

Frost CJ, Mescher MC, Dervinis C, Davis JM, Carlson JE, de Moraes CM. 2008. Priming defense genes and metabolites in hybrid poplar by the green leaf volatile cis-3-hexenyl acetate. New Phytologist 180: 722–734.

Fu J, Ren F, Lu X, Mao H, Xu M, Degenhardt J, Peters RJ, Wang Q. 2016. A tandem array of ent-kaurene synthases in maize with roles in gibberellin and more specialized metabolism. Plant Physiology 170: 742–751.

Glauser G, Vallat A, Balmer D. 2014. Hormone profiling. *Methods in molecular biology (Clifton*, N.J.) 1062: 597–608.

Goning M, Lopez-hilfiker F, Hutterli M, Erb M, Pfander M. 2022. Autosampler. Patent number US11227755B2. https://worldwide.espacenet.com/patent/search/family/061157140/publication/US11227755B2?q=erb%20tofwerk

González CV, Ibarra SE, Piccoli PN, Botto JF, Boccalandro HE. 2012. Phytochrome B increases drought tolerance by enhancing ABA sensitivity in Arabidopsis thaliana. Plant, Cell & Environment 35: 1958–1968.

Gouinguené SP, Turlings TCJ. 2002. The effects of abiotic factors on induced volatile emissions in corn plants. Plant Physiology 129: 1296–1307.

Helms AM, Moraes CM de, Tröger A, Alborn HT, Francke W, Tooker JF, Mescher MC. 2017. Identification of an insect-produced olfactory cue that primes plant defenses. Nature Communications 8: 337.

Holmes MG, Sager JC, Klein WH. 1986. Sensitivity to far-red radiation in stomata of Phaseolus vulgaris L.: Rhythmic effects on conductance and photosynthesis. Planta 168: 516–522.

Hu L. 2022. Integration of multiple volatile cues into plant defense responses. New Phytologist 233: 618–623.

Hu L, Ye M, Erb M. 2019. Integration of two herbivore-induced plant volatiles results in synergistic effects on plant defence and resistance. *Plant*, Cell & Environment 42: 959–971.

Johnson GN. 2011. Physiology of PSI cyclic electron transport in higher plants. Biochimica et biophysica acta 1807: 384–389.

Kegge W, Ninkovic V, Glinwood R, Welschen RAM, Voesenek LACJ, Pierik R. 2015. Red:far-red light conditions affect the emission of volatile organic compounds from barley (Hordeum vulgare), leading to altered biomass allocation in neighbouring plants. Annals of botany 115: 961–970.

Kegge W, Weldegergis BT, Soler R, Eijk MV-V, Dicke M, Voesenek LACJ, Pierik R. 2013. Canopy light cues affect emission of constitutive and methyl jasmonate-induced volatile organic compounds in Arabidopsis thaliana. New Phytologist 200: 861–874.

Kessler A, Baldwin IT. 2001. Defensive function of herbivore-induced plant volatile emissions in nature. *Science (New York*, N.Y.) 291: 2141–2144.

Kroes A, van Loon JJA, Dicke M. 2015. Density-dependent interference of aphids with caterpillar-induced defenses in Arabidopsis: involvement of phytohormones and transcription factors. Plant & cell physiology 56: 98–106.

Küpers JJ, Oskam L, Pierik R. 2020. Photoreceptors Regulate Plant Developmental Plasticity through Auxin. Plants 9: 940.

Leone M, Keller MM, Cerrudo I, Ballaré CL. 2014. To grow or defend? Low red: far-red ratios reduce jasmonate sensitivity in Arabidopsis seedlings by promoting DELLA degradation and increasing JAZ10 stability. New Phytologist 204: 355–367.

Lim J, Park J-H, Jung S, Hwang D, Nam HG, Hong S. 2018. Antagonistic Roles of PhyA and PhyB in Far-Red Light-Dependent Leaf Senescence in Arabidopsis thaliana. Plant & cell physiology 59: 1753–1764.

Lin P-A, Chen Y, Ponce G, Acevedo FE, Lynch JP, Anderson CT, Ali JG, Felton GW. 2022. Stomata-mediated interactions between plants, herbivores, and the environment. Trends in plant science 27: 287–300.

Maag D, Dalvit C, Thevenet D, Köhler A, Wouters FC, Vassão DG, Gershenzon J, Wolfender J-L, Turlings TCJ, Erb M et al. 2014. 3-β-D-Glucopyranosyl-6-methoxy-2-benzoxazolinone (MBOA-N-Glc) is an insect detoxification product of maize 1,4-benzoxazin-3-ones. Phytochemistry 102: 97– 105.

Maddonni GA, Otegui ME, Andrieu B, Chelle M, Casal JJ. 2002. Maize leaves turn away from neighbors. Plant Physiology 130: 1181–1189.

Martínez-García JF, Gallemí M, Molina-Contreras MJ, Llorente B, Bevilaqua MRR, Quail PH. 2014. The shade avoidance syndrome in Arabidopsis: the antagonistic role of phytochrome a and B differentiates vegetation proximity and canopy shade. PLOS ONE 9: e109275.

Meents AK, Mithöfer A. 2020. Plant-Plant Communication: Is There a Role for Volatile Damage-Associated Molecular Patterns? Frontiers in plant science 11: 583275.

Mérey GE von, Veyrat N, D’Alessandro M, Turlings TCJ. 2013. Herbivore-induced maize leaf volatiles affect attraction and feeding behavior of Spodoptera littoralis caterpillars. Frontiers in Plant Science 4: 209.

Mérey G von, Veyrat N, Mahuku G, Valdez RL, Turlings TCJ, D’Alessandro M. 2011. Dispensing synthetic green leaf volatiles in maize fields increases the release of sesquiterpenes by the plants, but has little effect on the attraction of pest and beneficial insects. Phytochemistry 72: 1838–1847.

Meza-Canales ID, Meldau S, Zavala JA, Baldwin IT. 2017. Herbivore perception decreases photosynthetic carbon assimilation and reduces stomatal conductance by engaging 12-oxo-phytodienoic acid, mitogen-activated protein kinase 4 and cytokinin perception. Plant, Cell & Environment 40: 1039–1056.

Moraes CM de, Mescher MC, Tumlinson JH. 2001. Caterpillar-induced nocturnal plant volatiles repel conspecific females. Nature 410: 577–580.

Moreira X, Nell CS, Katsanis A, Rasmann S, Mooney KA. 2018. Herbivore specificity and the chemical basis of plant-plant communication in Baccharis salicifolia (Asteraceae). New Phytologist 220: 703–713.

Moreno JE, Tao Y, Chory J, Ballaré CL. 2009. Ecological modulation of plant defense via phytochrome control of jasmonate sensitivity. Proceedings of the National Academy of Sciences of the United States of America 106: 4935–4940.

Mujiono K, Tohi T, Sobhy IS, Hojo Y, Shinya T, Galis I. 2021. Herbivore-induced and constitutive volatiles are controlled by different oxylipin-dependent mechanisms in rice. Plant, Cell & Environment 44: 2687–2699.

Nguyen D, Rieu I, Mariani C, van Dam NM. 2016. How plants handle multiple stresses: hormonal interactions underlying responses to abiotic stress and insect herbivory. Plant Molecular Biology 91: 727–740.

Niinemets U, Loreto F, Reichstein M. 2004. Physiological and physicochemical controls on foliar volatile organic compound emissions. Trends in plant science 9: 180–186.

Niinemets Ü. 2003. Controls on the emission of plant volatiles through stomata: Differential sensitivity of emission rates to stomatal closure explained. Journal of Geophysical Research 108.

Panda D, Dash GK, Mohanty S, Sekhar S, Roy A, Tudu C, Behera L, Tripathy BC, Baig MJ, 2023. Phytochrome A mediated modulation of photosynthesis, development and yield in rice (*Oryza sativa* L.) in fluctuating light environment. Environmental and Experimental Botany, 206, p.105183.

Paudel Timilsena B, Seidl-Adams I, Tumlinson JH. 2020. Herbivore-specific plant volatiles prime neighboring plants for nonspecific defense responses. Plant, Cell & Environment 43: 787–800.

Pierik R, Ballaré CL. 2021. Control of Plant Growth and Defense by Photoreceptors: From Mechanisms to Opportunities in Agriculture. Molecular Plant 14: 61–76.

Pierik R, Djakovic-Petrovic T, Keuskamp DH, Wit M de, Voesenek LACJ. 2009. Auxin and ethylene regulate elongation responses to neighbor proximity signals independent of gibberellin and della proteins in Arabidopsis. Plant Physiology 149: 1701–1712.

Pierik R, Wit M de. 2014. Shade avoidance: phytochrome signalling and other aboveground neighbour detection cues. Journal of experimental botany 65: 2815–2824.

Richter A, Schaff C, Zhang Z, Lipka AE, Tian F, Köllner TG, Schnee C, Preiß S, Irmisch S, Jander G et al. 2016. Characterization of Biosynthetic Pathways for the Production of the Volatile Homoterpenes DMNT and TMTT in Zea mays. The Plant cell 28: 2651–2665.

Richter A, Seidl-Adams I, Köllner TG, Schaff C, Tumlinson JH, Degenhardt J. 2015. A small, differentially regulated family of farnesyl diphosphate synthases in maize (Zea mays) provides farnesyl diphosphate for the biosynthesis of herbivore-induced sesquiterpenes. Planta 241: 1351– 1361.

Savchenko T, Kolla VA, Wang CQ, Nasafi Z, Hicks DR, Phadungchob B, Chehab WE, Brandizzi F, Froehlich J, Dehesh K. 2014. Functional convergence of oxylipin and abscisic acid pathways controls stomatal closure in response to drought. Plant Physiology 164: 1151–1160.

Schmelz EA, Alborn HT, Banchio E, Tumlinson JH. 2003. Quantitative relationships between induced jasmonic acid levels and volatile emission in Zea mays during Spodoptera exigua herbivory. Planta 216: 665–673.

Schnee C, Köllner TG, Gershenzon J, Degenhardt J. 2002. The maize gene terpene synthase 1 encodes a sesquiterpene synthase catalyzing the formation of (E)-beta-farnesene, (E)-nerolidol, and (E,E)-farnesol after herbivore damage. Plant Physiology 130: 2049–2060.

Schnee C, Köllner TG, Held M, Turlings TCJ, Gershenzon J, Degenhardt J. 2006. The products of a single maize sesquiterpene synthase form a volatile defense signal that attracts natural enemies of maize herbivores. Proceedings of the National Academy of Sciences of the United States of America 103: 1129–1134.

Seidl-Adams I, Richter A, Boomer KB, Yoshinaga N, Degenhardt J, Tumlinson JH. 2015. Emission of herbivore elicitor-induced sesquiterpenes is regulated by stomatal aperture in maize (Zea mays) seedlings. Plant, Cell & Environment 38: 23–34.

Sheehan MJ, Kennedy LM, Costich DE, Brutnell TP. 2007. Subfunctionalization of PhyB1 and PhyB2 in the control of seedling and mature plant traits in maize. The Plant journal: for cell and molecular biology 49: 338–353.

Soler R, Erb M, Kaplan I. 2013. Long distance root-shoot signalling in plant-insect community interactions. Trends in plant science 18: 149–156.

Ton J, D’Alessandro M, Jourdie V, Jakab G, Karlen D, Held M, Mauch-Mani B, Turlings TCJ. 2007. Priming by airborne signals boosts direct and indirect resistance in maize. The Plant journal: for cell and molecular biology 49: 16–26.

Vries J de, Poelman EH, Anten N, Evers JB. 2018. Elucidating the interaction between light competition and herbivore feeding patterns using functional-structural plant modelling. Annals of botany 121: 1019–1031.

Way DA, Pearcy RW. 2012. Sunflecks in trees and forests: from photosynthetic physiology to global change biology. Tree physiology 32: 1066–1081.

Wit M de, Keuskamp DH, Bongers FJ, Hornitschek P, Gommers CMM, Reinen E, Martínez-Cerón C, Fankhauser C, Pierik R. 2016. Integration of Phytochrome and Cryptochrome Signals Determines Plant Growth during Competition for Light. Current biology: CB 26: 3320–3326.

Wit M de, Spoel SH, Sanchez-Perez GF, Gommers CMM, Pieterse CMJ, Voesenek LACJ, Pierik R. 2013. Perception of low red:far-red ratio compromises both salicylic acid-and jasmonic acid-dependent pathogen defences in Arabidopsis. The Plant Journal 75: 90–103.

Xu G, Cao J, Wang X, Chen Q, Jin W, Li Z, Tian F. 2019. Evolutionary Metabolomics Identifies Substantial Metabolic Divergence between Maize and Its Wild Ancestor, Teosinte. The Plant Cell 31: 1990–2009.

Xue J, Gou L, Zhao Y, Yao M, Yao H, Tian J, Zhang W. 2016. Effects of light intensity within the canopy on maize lodging. Field Crops Research 188: 133–141.

Yang C, Xie F, Jiang Y, Li Z, Huang X, Li L. 2018. Phytochrome A Negatively Regulates the Shade Avoidance Response by Increasing Auxin/Indole Acidic Acid Protein Stability. Developmental Cell 44: 29–41.e4.

Ye M, Glauser G, Lou Y, Erb M, Hu L. 2019. Molecular Dissection of Early Defense Signaling Underlying Volatile-Mediated Defense Regulation and Herbivore Resistance in Rice. The Plant Cell 31: 687–698.

Zhang P-J, Broekgaarden C, Zheng S-J, Snoeren TAL, van Loon JJA, Gols R, Dicke M. 2013. Jasmonate and ethylene signaling mediate whitefly-induced interference with indirect plant defense in Arabidopsis thaliana. New Phytologist 197: 1291–1299.

Zhen S, Bugbee B. 2020. Far-red photons have equivalent efficiency to traditional photosynthetic photons: Implications for redefining photosynthetically active radiation. Plant, Cell & Environment 43: 1259–1272.

Zhen S, van Iersel MW, Bugbee B. 2022. Photosynthesis in sun and shade: the surprising importance of far-red photons. New Phytologist. doi: 10.1111/nph.18375.

