## Supplemental information for "Far-red light increases maize volatile emissions in response to volatile cues from neighboring plants"

**Table S1.** RT-qPCR primers

| *Target gene* | *Gene name* | *Primers* | *Reference* |
|---|---|---|---|
| *ZmActin* | *Zea mays*<br>*Actin* | CCATGAGGCCACGTACAACT<br>GGTAAAACCCCCACTGAGGA | Erb *et al.*, 2009 |
| *ZmFPPS3* | *Zea mays*<br>*Farnesyl pyrophosphate synthase 3* | CCTGGCTAGTTGTGCAAGCT<br>GAAAACAGTTTGGACTGCCT | Seidl-Adams *et al.*, 2015 |
| *ZmTPS1* | *Zea mays*<br>*Terpene synthase 1* | GGATAGCATAGGCTCAAGGT<br>GAAGGGACCGTCCAGAGCAT | Xu *et al.*, 2019 |
| *ZmTPS2* | *Zea mays*<br>*Terpene synthase 2* | TACCGGGTCGAGATCACCAA<br>TCGTTCGTAACGGTGTGGAG | n.a. |
| *ZmTPS10* | *Zea mays*<br>*Terpene synthase 10* | TGTGTCCACGGTCCAATGTT<br>GTCCGCTGTCCTTGCAAAAT | Schnee *et al.*, 2006 |
| *ZmCYP92C5* | *Zea mays*<br>*P450 monooxygenase* | ACGACCTTCACGACCATTTC<br>CCTCATCCAGGACATCATCG | Richter *et al.*, 2016 |

**Supplemental Method S1**

Phytohormones were extracted with ethyl acetate: formic acid, 99.5:0.5 (v/v) spiked with isotopically labelled standards (1 ng of d5-JA, 13C6-JA-Ile, d4-SA, and d6-ABA) and analysed by ultra-high-performance liquid chromatography tandem mass spectrometry (UHPLC-MS/MS). Approximately 100 mg of frozen and homogenized leaf material was transferred to a 1.5-ml Eppendorf tubes and extracted with 1 ml of ethyl acetate: formic acid, 99.5:0.5 (v/v) spiked with isotopically labelled standards (1 ng of d5-JA, 13C6-JA-Ile, d4-SA, and d6-ABA). Samples were then vortexed for 10 s and extracted in a mixer mill at a frequency of of 30 Hz for 3 min. Next, samples were centrifuged at 14000 g for 3 min at 4 °C. The supernatants were transferred to new Eppendorf tubes and the pellets were re-extracted with 0.5 mL of ethyl acetate: formic acid, 99.5:0.5 (v/v) and centrifuged at 14000 g for 3 min at 4 °C. The two supernatants were combined and evaporated to dryness. Residues were re-suspended in 200 µL of MeOH 50 %. The suspensions were transferred to 0.2 mL microcentrifuge tubes and centrifuged for 1.5 min at 14,000 × *g*. Supernatants were transferred to GC-MS vials for analyses. The final concentrations of internal standards were 5 ng/mL for d5-JA, d6-ABA, d6-SA, 13C6-JA-Ile and d5-IAA. Two microliters of the extracts were analyzed using an Acquity UPLC (Waters) coupled to a QTRAP 6500+ (Sciex) following the protocol described in (Glauser *et al.*, 2014)

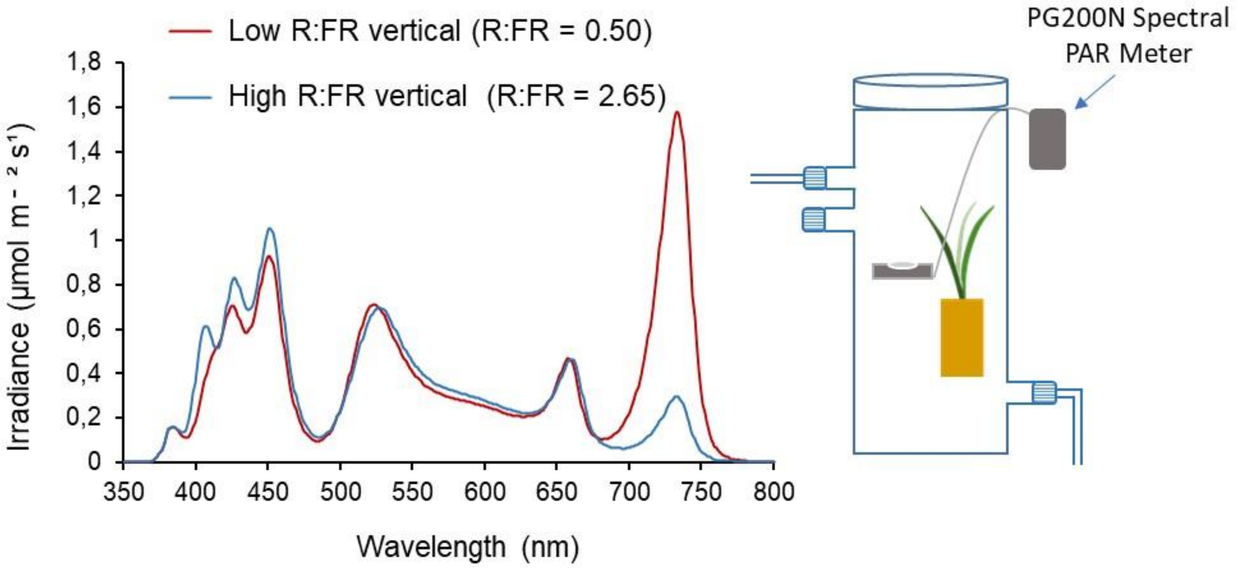


**Figure S1. Light spectral composition determined in far-red (FR) supplementation experiments using a PG200N Spectral PAR Meter (UPRtek).** Light spectra were vertically measured inside two representative glass chambers where 11-day-old maize (*Zea mays*) plants were exposed to low or high R:FR light conditions. Photosynthetically active radiation values under high or low R:FR light conditions in these two glass chambers were 121 and 111 µmol $m^{-2}$ $s^{1}$ respectively. Averaged PAR levels among experimental replicates for both low and high R:FR light treatments were the same, i.e., 120 ± 15 µmol $m^{-2}$ $s^{1}$. The legend indicates the ratio between Red (R: λ 600-700 nm) and Far Red (FR: λ 700-800 nm) light.

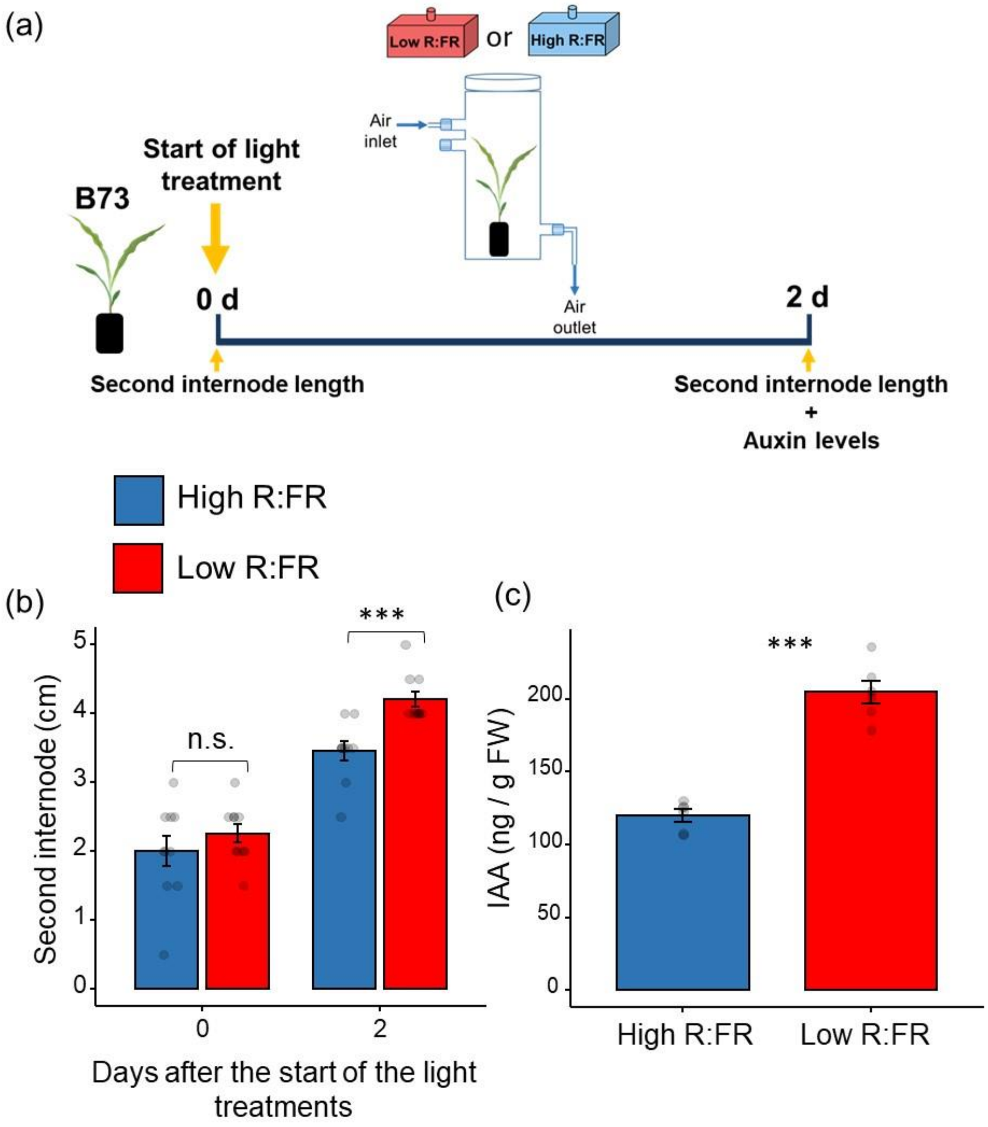


**Figure S2. Effects of far-red enrichment on maize growth and auxin levels. (a)** Schematic overview of the experimental set up. Eleven-day old `B73´ maize (*Zea mays*) plants were placed inside transparent glass chambers and exposed to low or high R:FR light conditions by modulating FR levels for 2 days. **(b)** Length of the second internode (mean ± SEM, $n$ = 10) measured before and after 2 d of light treatments. **(c)** Auxin (IAA) concentrations (mean ± SEM, $n$ = 12) measured after 2 d of light treatments. Asterisks denote differences between treatments tested by Student *t*-tests. *** $p < 0.001$. n.s. not significant.

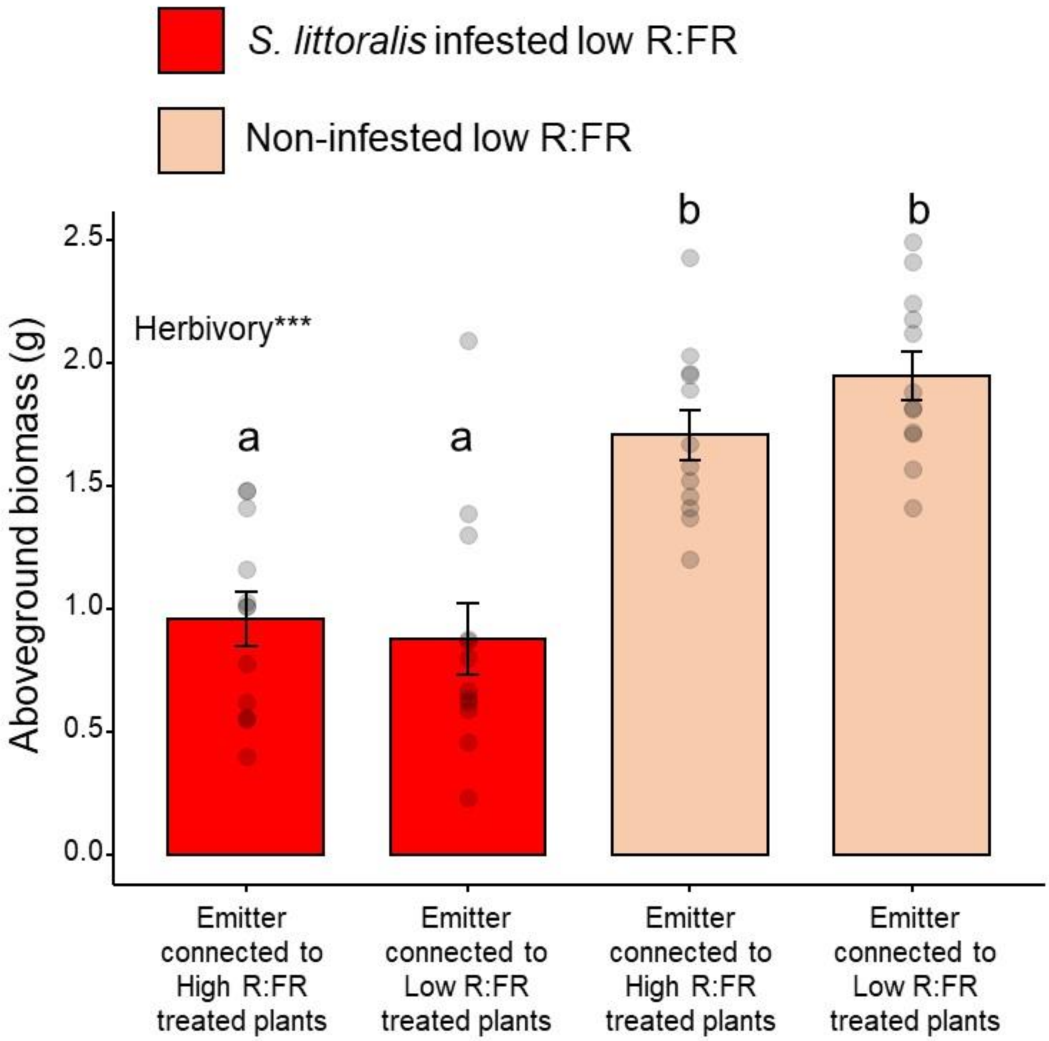


**Figure S3.** Aboveground biomass (mean ± SEM, $n$ = 12) of low R:FR treated maize plants that were either infested with ten second-instar *S. littoralis* larvae or left non-infested. Measurements were performed at 22 h after the start of the infestation's treatments. Effect of herbivory was tested by ANOVA. Different letters denote significant differences among treatments tested by Tukey HSD post hoc test at $p < 0.05$. *** $p < 0.001$.

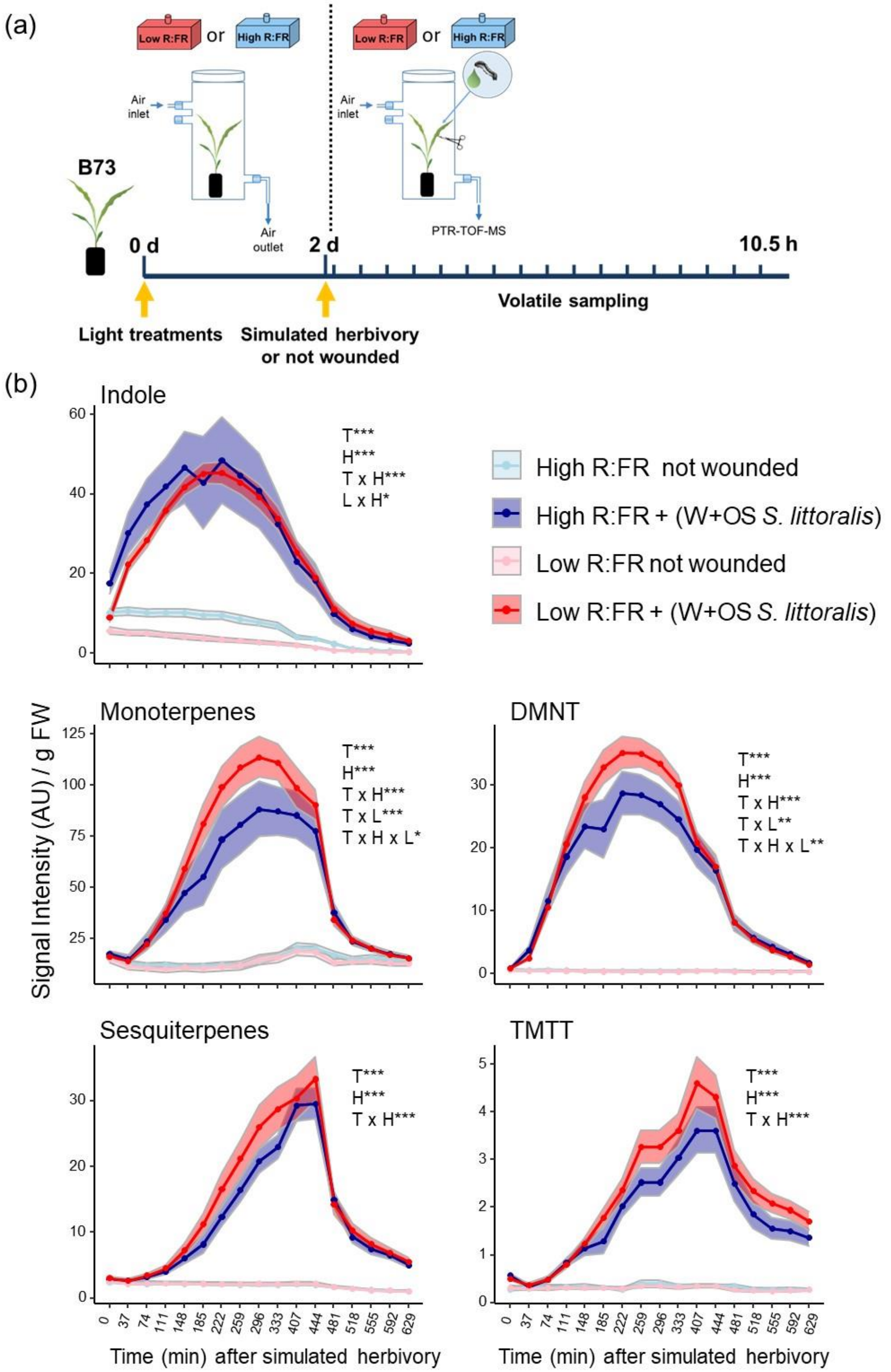


**Figure S4. Effect of far-red enrichment on herbivory-induced volatile emission in maize. (a)** Schematic overview of the experimental set-up. Eleven-day old `B73´ maize (*Zea mays*) plants were exposed to low or high R:FR conditions for 2 d by modulating FR levels, and subsequently induced with simulated herbivory (wounding and application of *Spodoptera littoralis* oral secretions; W+OS) or left unwounded. **(b)** Time series analysis of volatile emissions (mean ± SEM, *n* = 8) determined by PTR-TOF-MS at intervals of 37 min in high and low R:FR-treated B73 maize (*Zea mays*) plants after simulated herbivory. Volatile measurements started at 12:30 pm, 10 min after simulated herbivory treatment. The effects of sampling time (T), light treatment (L), simulated herbivory (H) and their interactions on volatile emission were tested using linear mixed-effects models. Plant unit was included as the random intercept as well as a correlation structure when autocorrelation among residuals was found significant ($p < 0.05$). Statistically significant effects are indicated in each graph. *$p < 0.05$, ** $p < 0.001$, *** $p < 0.001$. DMNT and TMTT stand for the homoterpenes (*E*)-4,8-dimethyl-1,3,7-nonatriene and (*E, E*)-4,8,12-trimethyltrideca-1,3,7,11-tetraene, respectively. AU refers to arbitrary units.

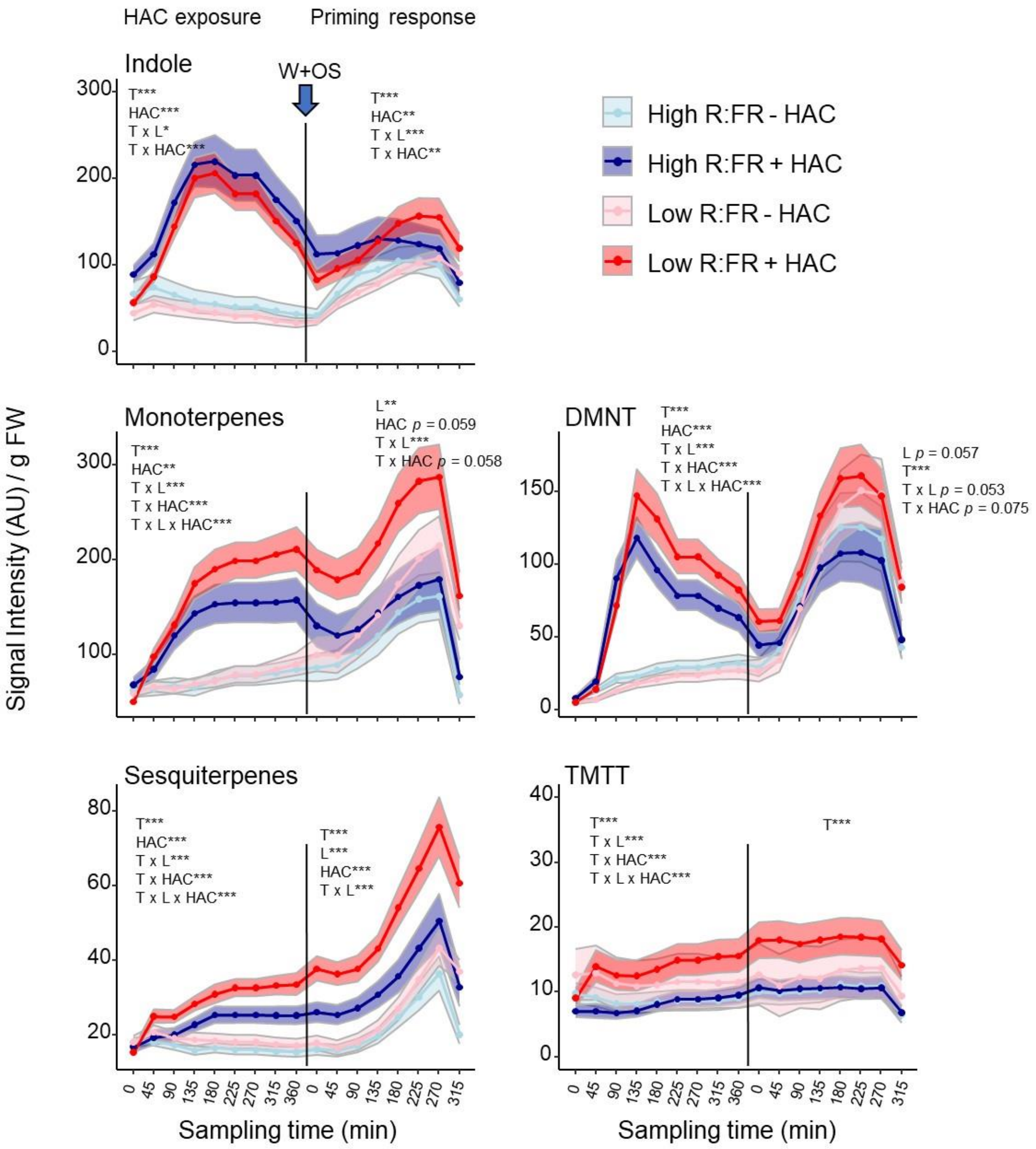


**Figure S5. Far-red enrichment boosts maize responses to (*Z*)-3-hexenyl acetate in 30-day old maize (*Zea mays*) plants.** Time-series analysis of volatile emissions (mean ± SEM, $n = 11$) determined by PTR-TOF-MS in low and high R:FR treated 30-day-old B73 plants during exposure to (*Z*)-3-hexenyl acetate (+ HAC) or clean air (- HAC) and after induction with simulated herbivory (wounding and application of *Spodoptera littoralis* oral secretions; W+OS) (priming response). HAC exposure and volatile measurements started between 9:00 and 10:00 am. The effects of the sampling time (T), HAC, light treatment (L) and their interactions on volatile emissions were tested separately for the HAC exposure and priming phase using linear mixed-effects models. Statistically significant effects are shown in each graph (*$p < 0.05$, ** $p < 0.001$, *** $p < 0.001$). DMNT and TMTT stand for the homoterpenes (*E*)-4,8-dimethyl-1,3,7-nonatriene and (*E*,*E*)-4,8,12-trimethyltrideca-1,3,7,11-tetraene, respectively. AU refers to arbitrary units.

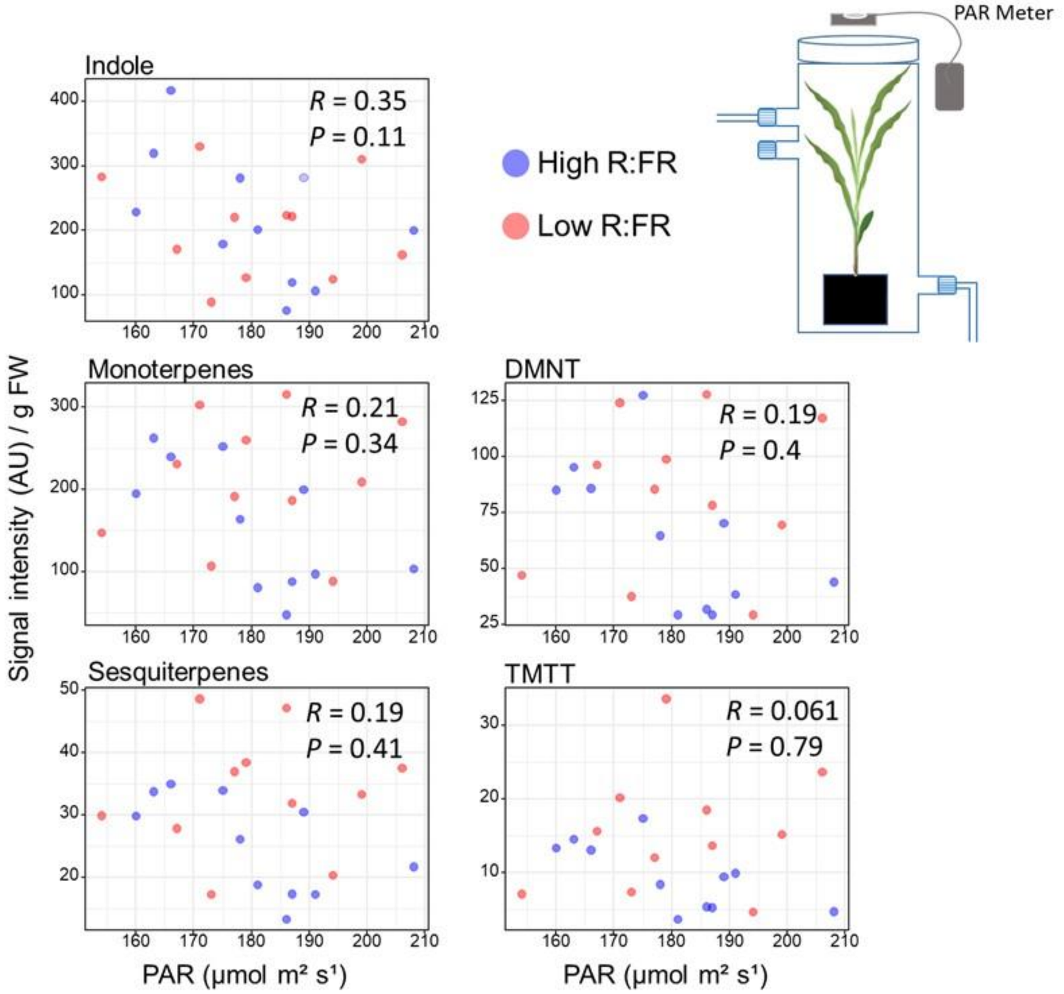


**Figure S6. Spatial variations in photosynthetically active radiation (PAR) levels in the experimental set up did not influence 30-day-old maize responses to (*Z*)-3-hexenyl acetate.** Scatter plots depict the relationship between the PAR level measured over the lid of the transparent glass chamber containing the test plant and its volatile emissions during (*Z*)-3-hexenyl acetate exposure. Signal intensity for monoterpenes, sesquiterpenes, linalool, DMNT and TMTT were obtained from the time points corresponding to 180, 360, 360, 360, 135, and 315 min, respectively, after the start of (*Z*)-3-hexenyl acetate exposure treatment displayed in Supplemental Fig. S6. The Pearson correlation coefficient (*R*) and *P* value are indicated. DMNT and TMTT stand for the homoterpenes (*E*)-4,8-dimethyl-1,3,7-nonatriene and (*E*,*E*)-4,8,12-trimethyltrideca-1,3,7,11-tetraene, respectively. AU refers to arbitrary units.

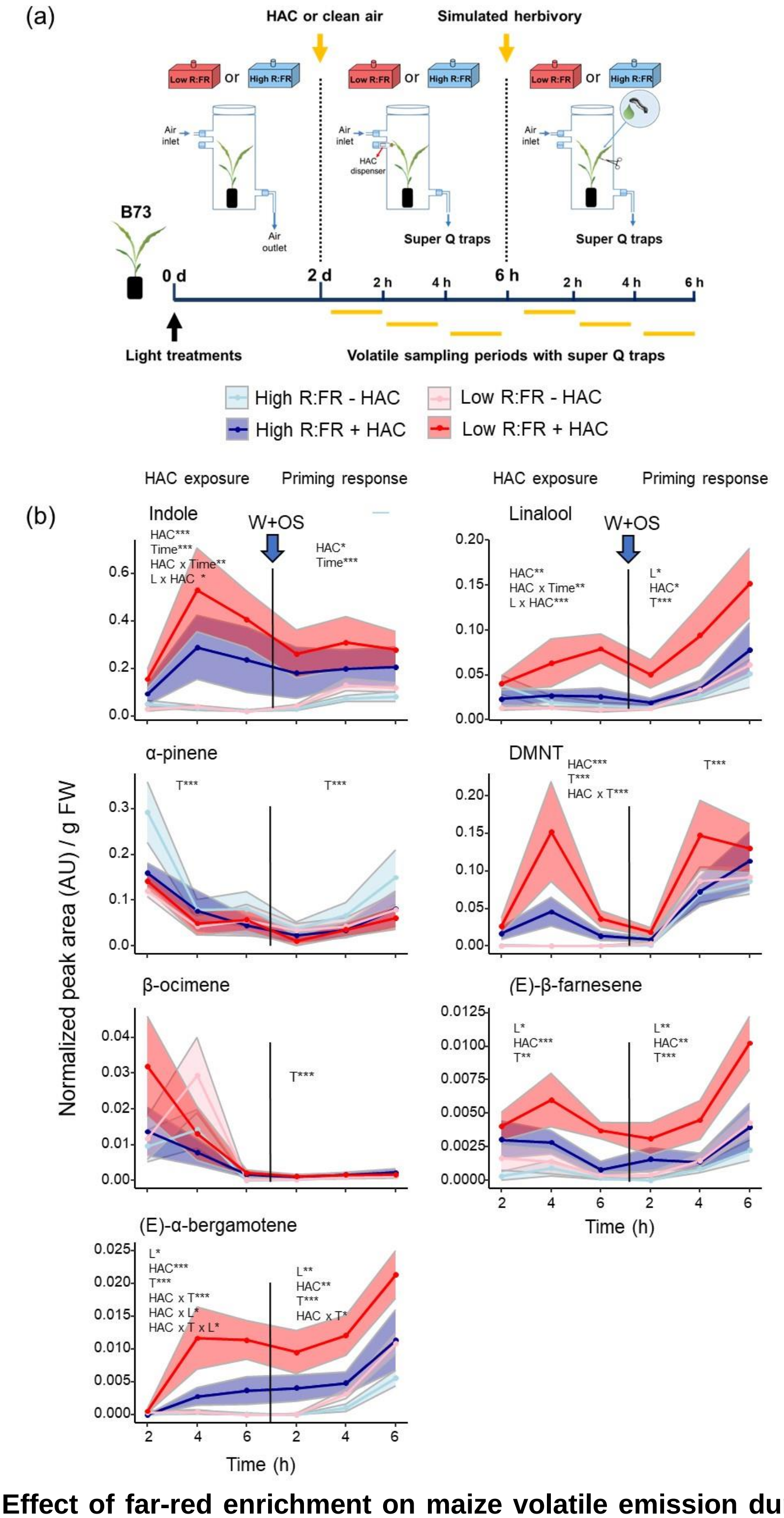


**Figure S7. Effect of far-red enrichment on maize volatile emission during exposure to (*Z*)-3-hexenyl acetate and after simulated herbivory. (a)** Schematic overview of the experimental set-up and sampling methods. Eleven-day old `B73´ maize (*Zea mays*) plants were exposed to low or high

R:FR conditions for 2 d by modulating FR levels, and subsequently exposed to (*Z*)-3-hexenyl acetate (HAC) or clean air for 6 h. After HAC treatment, all plants were induced with simulated herbivory (wounding and application of *Spodoptera littoralis* oral secretions). **(b)** Time-series analysis of volatiles emissions (mean ± SEM, $n = 6$) determined in low and high R:FR-treated B73 maize (*Zea mays*) plants during exposure to (*Z*)-3-hexenyl acetate (+ HAC) or clean air (- HAC) for 6 h, and after induction with simulated herbivory (priming response). +/- HAC treatments and volatile collection started at 9:00 am. Volatiles were then collected for periods of 2 h for 12 h using superQ traps. Eluded extracts were analysed by GC/MS. The effects of the sampling time (T), HAC, light treatment (L) and their interactions on volatiles emission were tested using linear mixed-effects models. Statistically significant effects are shown in each graph. *$p < 0.05$, ** $p < 0.001$, *** $p < 0.001$. DMNT stands for the homoterpene (*E*)-4,8-dimethyl-1,3,7-nonatriene. AU refers to arbitrary units.

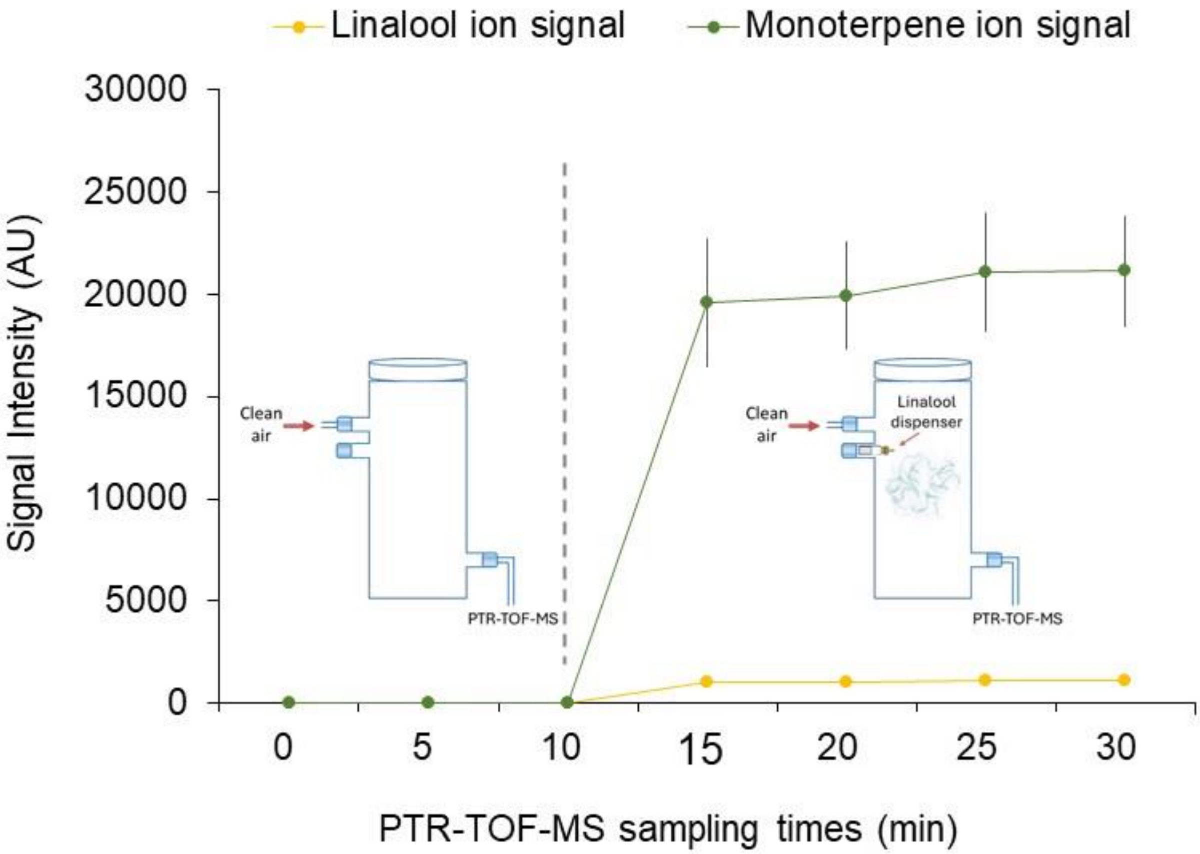


**Figure S8. PTR-TOF-MS analysis of pure linalool.** Time-series measurements (n = 5, ± SEM) of the linalool signal (*m*/*z* 137) and the monoterpenes signal (*m*/*z* 137) from dispensers containing 100 µL of pure linalool (SIGMA). Emission of the linalool dispensers were detected using PTR-TOF-MS at intervals of 60 seconds.

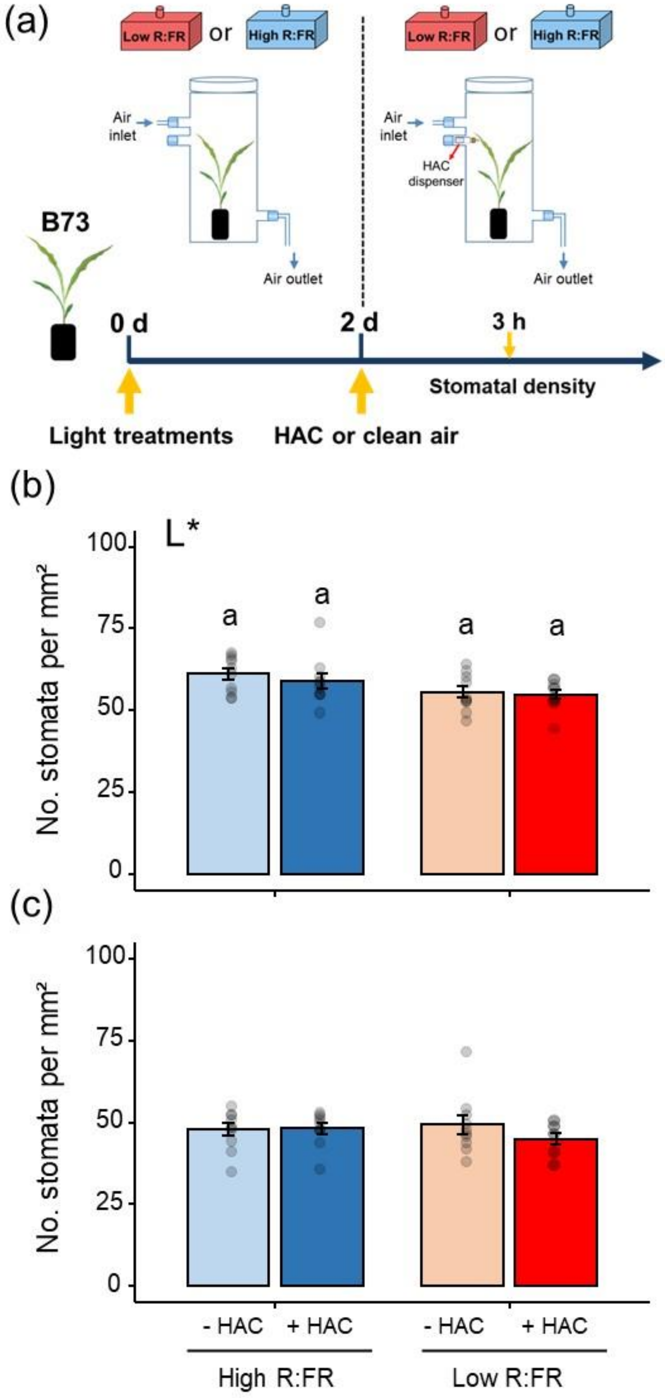


**Figure S9. Effect of far-red enrichment and (*Z*)-3-hexenyl acetate (HAC) on stomatal density. (a)** Schematical overview of the experimental set up and sampling time. Density of stomata (mean ± SEM, $n$ = 10 individual plants; 8-10 pictures per plant and leaf) determined in the abaxial side of **(b)** leaf 3 and **(c)** leaf 2 from the bottom of high and low R:FR-treated B73 maize plants at 3 h after the start of HAC exposure. Effects of light treatment (L) and HAC on stomatal density were tested by two-way ANOVAs. Statistically significant effects are shown in the graphs. $*p < 0.05$. Different letters denote significant differences among groups tested by Tukey-HSD post hoc test at $p < 0.05$.

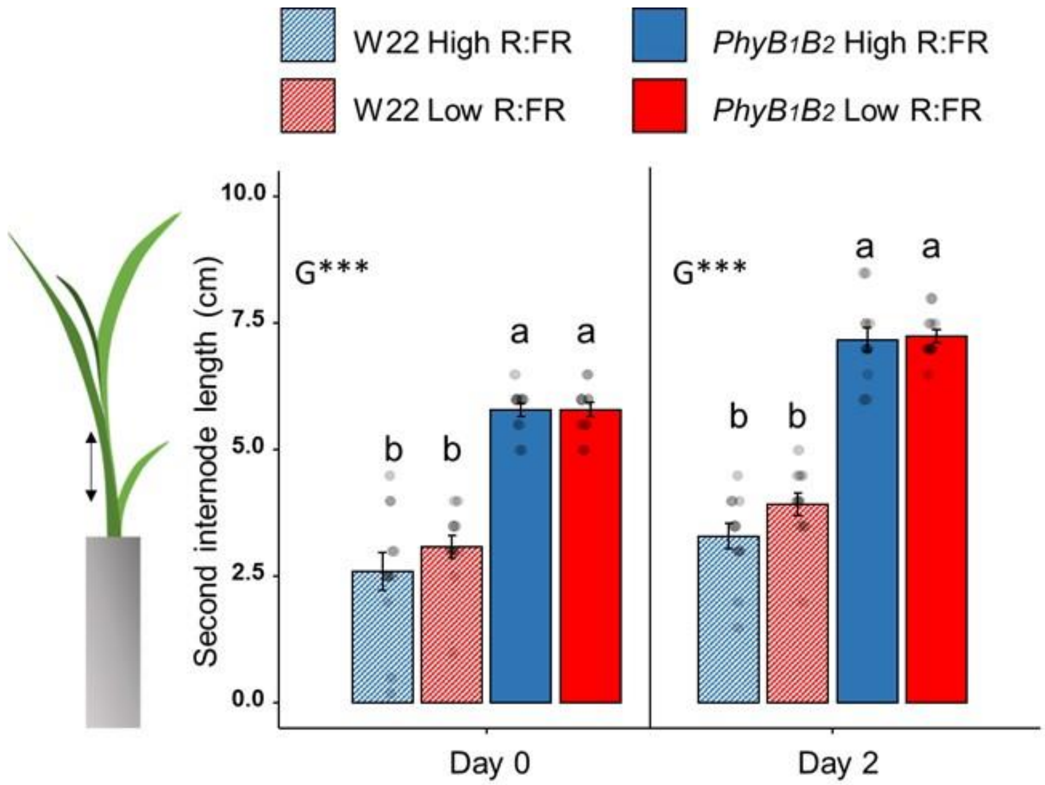


**Figure S10. Effect of far-red enrichment on the *phyB1phyB2* maize double mutant growth.** Length of the second internode (mean ± SEM, $n = 12$) was measured in *phyB1phyB* double mutant maize (*Zea mays*) plants and its wild type (inbred line `W22´) before and two days after the start of low and high R:FR light conditions. Effects of light, plant genotype and their interaction on second internode length were tested by two-way ANOVAs at the two time points. Statistically significant effects are shown in the graph with *$p < 0.05$, ** $p < 0.001$, *** $p < 0.001$. Different letters denote significant differences among groups tested by Tukey-HSD post hoc test at $p < 0.05$.

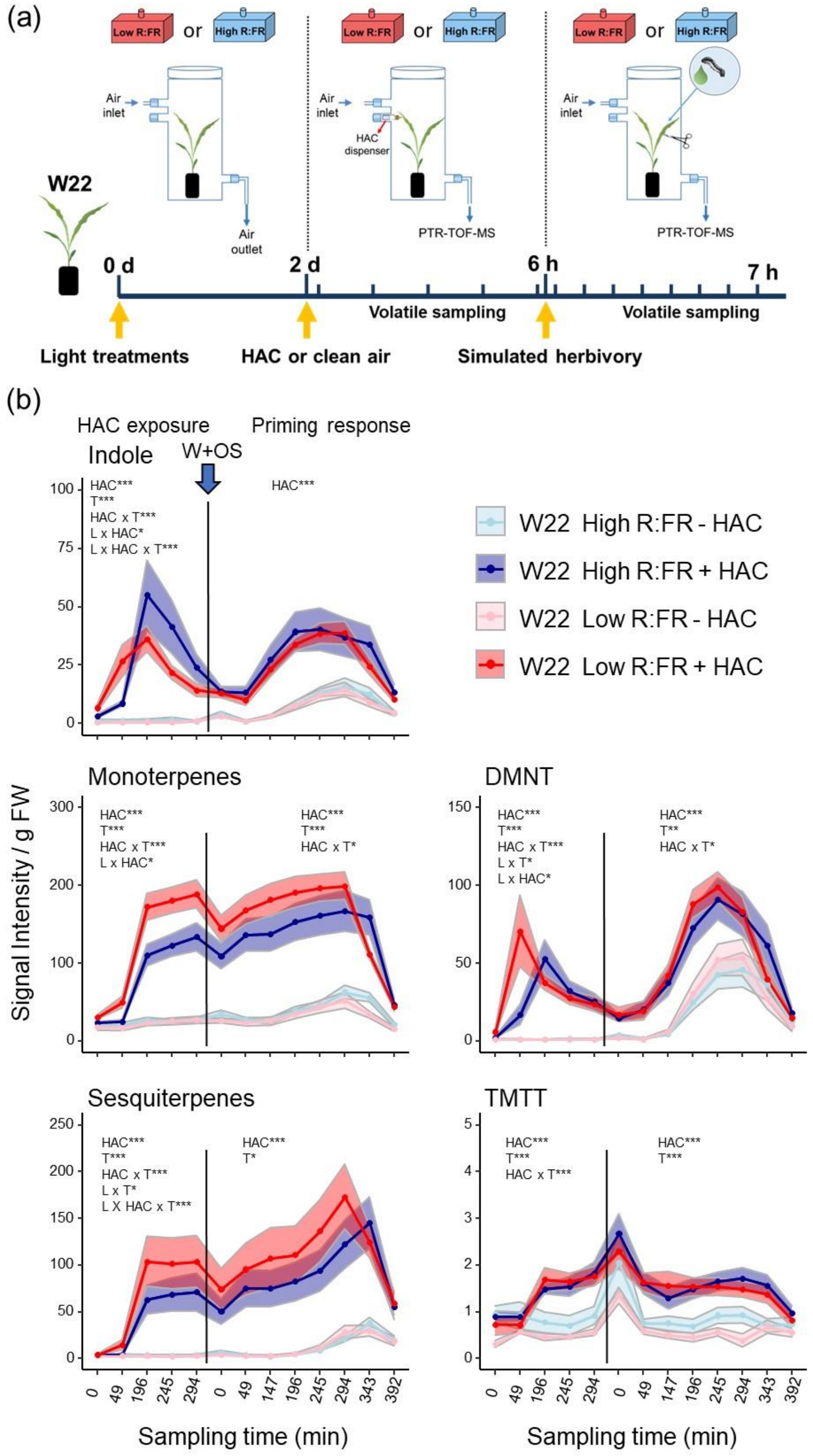


**Figure S11. Effect of far-red enrichment on volatiles emission during exposure to (*Z*)-3-hexenyl acetate and after simulated herbivory in the maize inbred line W22. (a)** Schematic overview of the experimental set up and sampling times. **(b)** Time-series analysis of volatiles emissions (mean ± SEM, *n* = 6) determined in low and high R:FR-treated \`W22´maize (*Zea* mays) plants by PTR-TOF-MS during exposure to (*Z*)-3-hexenyl acetate (+ HAC) or clean air (- HAC) and after induction with simulated herbivory (wounding and application of *Spodoptera littoralis* oral secretions; W+OS) (priming response). Volatile emissions were measured every 63 min, with +/- HAC treatments starting between 9 and 10:00 am. The effects of the sampling time (T), HAC, light treatment (L) and their interactions on HIPVs emission were tested using linear mixed effects models. Statistically significant effects are shown in each graph. *$p < 0.05$, ** $p < 0.001$, *** $p < 0.001$. DMNT and TMTT stand for the homoterpenes (*E*)-4,8-dimethyl-1,3,7-nonatriene and (*E, E*)-4,8,12-trimethyltrideca-1,3,7,11-tetraene, respectively. AU refers to arbitrary units.

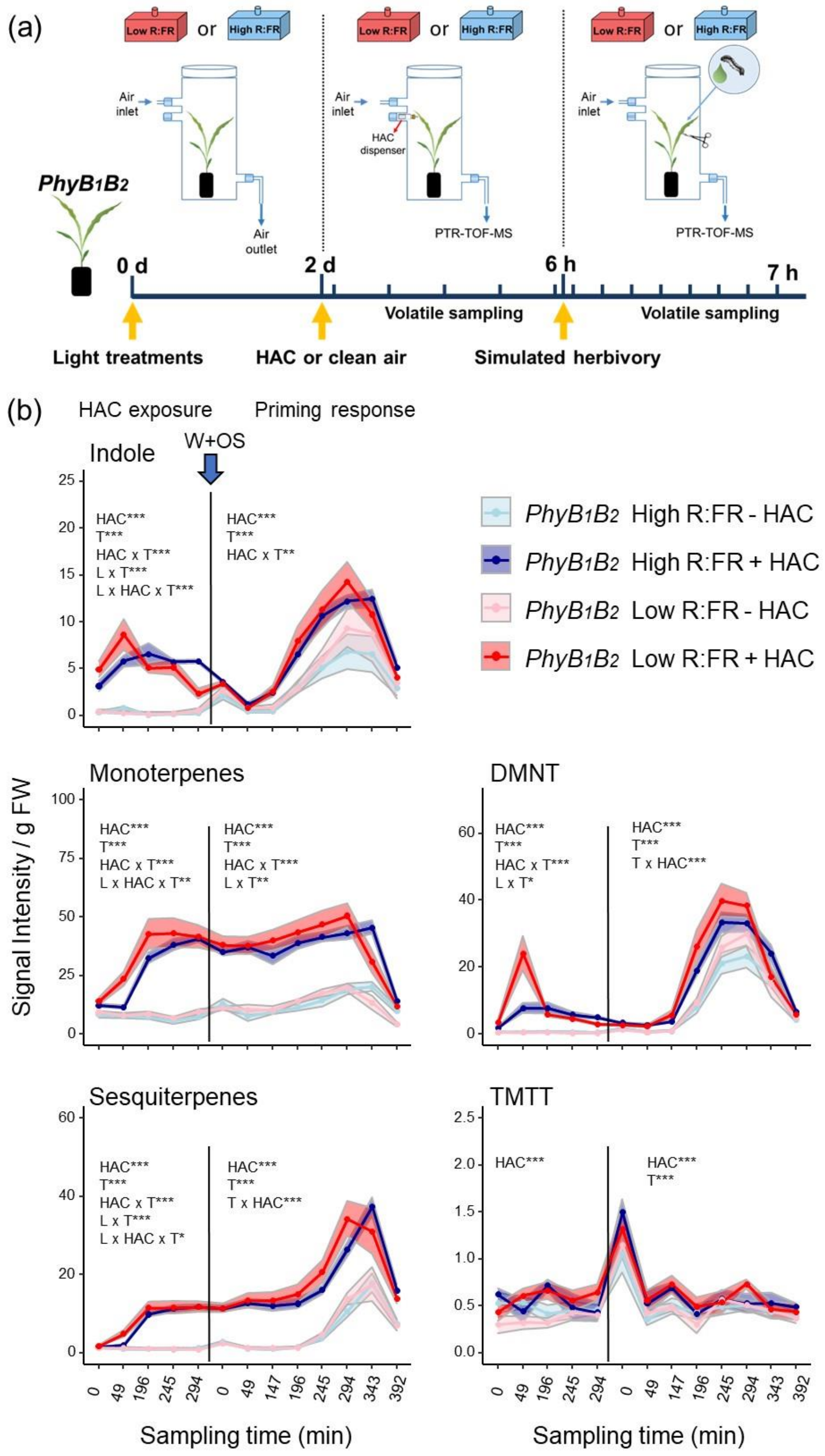


**Figure S12. Effect of far-red enrichment on volatiles emission during exposure to (*Z*)-3-hexenyl acetate and after simulated herbivory in the *phyB1phyB2* maize double mutant. (a)** Schematic overview of the experimental set up and sampling times. **(b)** Time-series analysis of volatiles emissions (mean ± SEM, *n* = 6) determined in low and high R:FR-treated *phyB1phyB2* maize plants by PTR-TOF-MS during exposure to (*Z*)-3-hexenyl acetate (+ HAC) or clean air (- HAC) and after induction with simulated herbivory (wounding and application of *S. littoralis* oral secretions; W+OS) (priming response). Volatile emissions were measured every 63 min, with +/- HAC treatments starting between 9 and 10:00 am. The effects of the sampling time (T), HAC, light treatment (L) and their interactions on HIPVs emission were tested using linear mixed effects models. Statistically significant effects are shown in each graph. $^*p < 0.05$, $^{**}\ p < 0.001$, $^{***}\ p < 0.001$. DMNT and TMTT stand for the homoterpenes (*E*)-4,8-dimethyl-1,3,7-nonatriene and (*E, E*)-4,8,12-trimethyltrideca-1,3,7,11-tetraene, respectively. AU refers to arbitrary units.

**References**


**Christensen SA, Nemchenko A, Borrego E, Murray I, Sobhy IS, Bosak L, DeBlasio S, Erb M, Robert CAM, Vaughn KA *et al.* 2013.** The maize lipoxygenase, ZmLOX10, mediates green leaf volatile, jasmonate and herbivore-induced plant volatile production for defense against insect attack. *The Plant Journal* **74**: 59–73.

**Erb M, Flors V, Karlen D, Lange E** de, **Planchamp C, D'Alessandro M, Turlings TCJ, Ton J. 2009.** Signal signature of aboveground-induced resistance upon belowground herbivory in maize. *The Plant Journal* **59**: 292–302.

**Glauser G, Vallat A, Balmer D. 2014.** Hormone profiling. *Methods in molecular biology (Clifton, N.J.)* **1062**: 597–608.

**Richter A, Schaff C, Zhang Z, Lipka AE, Tian F, Köllner TG, Schnee C, Preiß S, Irmisch S, Jander G *et al.* 2016.** Characterization of Biosynthetic Pathways for the Production of the Volatile Homoterpenes DMNT and TMTT in Zea mays. *The Plant cell* **28**: 2651–2665.

**Schnee C, Köllner TG, Held M, Turlings TCJ, Gershenzon J, Degenhardt J. 2006.** The products of a single maize sesquiterpene synthase form a volatile defense signal that attracts natural enemies of maize herbivores. *Proceedings of the National Academy of Sciences of the United States of America* **103**: 1129–1134.

**Seidl-Adams I, Richter A, Boomer KB, Yoshinaga N, Degenhardt J, Tumlinson JH. 2015.** Emission of herbivore elicitor-induced sesquiterpenes is regulated by stomatal aperture in maize (Zea mays) seedlings. *Plant, Cell & Environment* **38**: 23–34.

**Xu G, Cao J, Wang X, Chen Q, Jin W, Li Z, Tian F. 2019.** Evolutionary Metabolomics Identifies Substantial Metabolic Divergence between Maize and Its Wild Ancestor, Teosinte. *The Plant Cell* **31**: 1990–2009.
